# Reversal of vein of Galen aneurysmal malformation by stimulation of flow-mediated vessel fusion

**DOI:** 10.1101/2024.10.18.618370

**Authors:** Edwige Martin-Valiente, Yao Du, Chloé Goemans, Egor Zindy, Myckel Adam, Benoît Scheid, Miikka Vikkula, Boris Lubicz, Benoit Vanhollebeke, Nicolas Baeyens

## Abstract

Congenital vascular malformations arise from defective homeostatic development of the vascular tree^1^. The aneurysmal malformation of the Vein of Galen (VGAM) is the most frequent neurovascular malformation in neonates, with limited therapeutic options and poor outcomes in the most severe cases^2^. This congenital disease is consecutive to germline genetic mutation of *RASA1* or *EPHB4*^3,4^, but little is known about the mechanisms leading to its development. We generated mutant *rasa1a* and *ephb4a* deficiency models in zebrafish reproducing the genetic and structural characteristics of the VGAM in the dorsal longitudinal vein of the cerebral vasculature. We link the development of the malformation to a failure of the fusion of precursor blood vessels into a draining vessel for the choroidal type malformations and to a failure to constrict for the mural type malformations. The fusion process is driven by blood flow, sensed and integrated by endothelial cells. RASA1 deficiency destabilizes the homeostatic response to blood flow and contributes to impaired flow-mediated activation of MAPK and PI3K signaling. We targeted these defective mechanotransduction mechanisms pharmacologically in both *rasa1a* and *ephb4a* mutant models, successfully reestablishing the fusion and constriction processes in preexisting malformations. This work identifies molecular actors of the flow-mediated blood vessel fusion mechanism, a specific angiogenetic program, and provides ground for treating VGAM and other vascular remodeling disorders.

## Introduction

The formation of the vascular network involves a finely regulated chain of events, from generating polarized tubular structures forming a lumen to blood vessel specification. This network further consolidates in a mature tree, with arterioles pouring blood into the capillary network, then collecting back in venules and veins, providing optimal perfusion of the organs. The maturation process requires remodeling by pruning^5^, intussusception^6^, or fusion^7^ of immature vessels during development. These remodeling events happen in perfused vessels and are directly controlled by endothelial cells’ perception of flowing blood, with mechanotransduction acting as a critical regulator in the maturation of the vascular network^1,8^. Congenital vascular anomalies are morphogenetic defects of the vascular system, where the vascular tree’s hierarchization and the vessels’ architecture are locally affected.

Congenital vascular anomalies, especially high-flow arteriovenous malformations (AVMs), can develop during the flow-dependent maturation of the vascular tree. High-flow AVMs associated with loss of function mutations of bone morphogenic (BMP) signaling, as seen in hereditary hemorrhagic telangiectasia (HHT), are consecutive to disturbed vessel homeostasis. Disturbed endothelial cell quiescence mediated by fluid forces generated by blood flow leads to arteriovenous shunts in high-shear stress areas^9,10^. Sensing and integrating physiological values of laminar blood flow are associated with anti-inflammatory and stabilization programs essential for vessel homeostasis. BMP signaling is associated with a physiological shear stress set point^11^, underlining its importance in flow-associated vessel quiescence. An inadequate balance in flow-mediated BMP signaling leads to endothelial cell proliferation and migration, improper recruitment of mural cells, and the formation of large, fragile, and prone-to-rupture arteriovenous shunts^9^. We hypothesized that different flow transduction signaling programs might be perturbed as congenital vascular anomalies are not restricted to BMP-related genetic mutations and develop in various vascular beds subjected to specific fluid forces. Therefore, genetic disturbance of different mechanosensitive pathways key would lead to different vascular defects in the vascular tree^12,13^.

Vein of Galen Aneurysmal Malformations (VGAMs) are congenital arteriovenous fistulas draining to the median vein of the prosencephalon, or vein of Markowski, which is the vein of Galen embryonic precursor. Although rare, VGAMs are the most frequent cerebral arteriovenous shunts in neonates and infants^2^. Two major germline mutations have been reported in VGAM patients, with loss-of-function mutations in *RASA1* or *EPHB4* as the most prevalent lesions^3,4^. All the patients are heterozygous autosomal dominant carriers and often display other vascular malformations, regrouped in a series of clinical manifestations: capillary malformation-arteriovenous malformations (CM-AVMs)^14^, the Parkes-Weber syndrome^15^ and other fistulas. Several somatic mutations of *RASA1* have been reported, mostly in cutaneous lesions developing late in life^16–19^. Embolization by the trans-arterial route is the first (and only) line of treatment for VGAM. In neonates, endovascular treatment is a high-risk procedure, and when possible, the child is medically treated for congestive heart failure, the main life-threatening complication. Untreated VGAMs have a poor prognosis and are almost always fatal, but it is equally important to recognize that surgical treatment is inappropriate in some cases^20–22^. If there is evidence of preexisting brain damage^23^ (progressive atrophy or “melting brain syndrome,” parenchymal calcification) or severe multiorgan failure, a poor outcome, death, or survival with severe brain damage is inevitable. Current clinical management of VGAMs has reached its limits, and there is an urgent need to understand the disease’s etiology and identify complementary and supportive treatments.

For all these reasons, we sought to develop an *in-vivo* model of VGAMs to characterize the cellular mechanism mediating its formation and determine the most appropriate pharmacological approach to stabilize or reduce VGAMs and CHF development in neonates. Previous works by other teams have identified vascular remodeling defects in zebrafish larvae injected with antisense morpholinos against either *rasa1a*^24^ or *ephb4a*^25^, loss of *ephb4a* leading to the formation of fistulae in the dorsal longitudinal vein (DLV), reminiscing the fistulae observed in the VGAM patients^25^.

### Development of a clinically relevant animal model of VGAMs

VGAMs are clinically classified into two main types. The first one, “choroidal”, presents multiple high-flow fistulas and is the most complex manifestation, with a poorer prognosis. The second one, “mural”, is generally a single fistula generating a dilated median vein ^26^ (Figure 1A).

**Figure 1:**
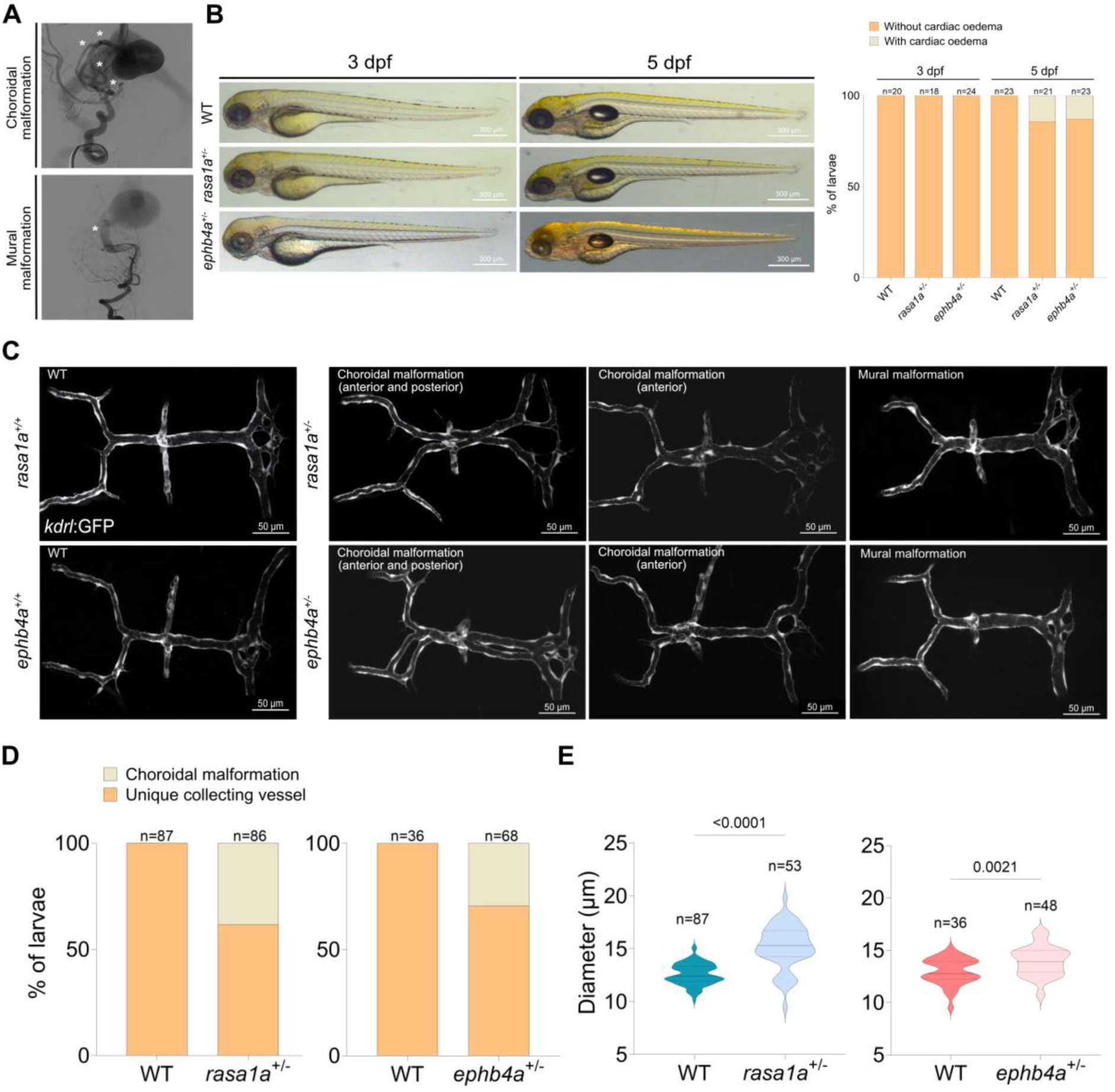
Development of *rasa1a and ephb4a* mutants as preclinical models of VGAMs. **A.** Digital subtraction angiography (Philips, Allura biplane system) frontal view of the right ICA and left VA showing a choroidal type VGAM in the upper panel and a mural type in the lower panel **B.** Lateral views of WT (*rasa1a^+/+^*or *ephb4a^+/+^* ), *rasa1a^+/-^* and *ephb4a^+/-^*larvae at 3 dpf and 5 dpf. Quantification of cardiac edema in WT, *rasa1a^+/-^* and *ephb4a^+/-^* larvae at 3 dpf (n=20 WT, n=18 *rasa1a^+/-^* and n=24 *ephb4a^+/-^*) and 5 dpf (n=23 WT, n=21 *rasa1a^+/-^* and n=23 *ephb4a^+/-^*). **C.** Dorsal view of the dorsal longitudinal vein (DLV) of WT (*rasa1a^+/+^* and *ephb4a^+/+^*), *rasa1a^+/-^* and *ephb4a^+/-^*larvae at 3 dpf. **D.** Quantification of the percentage of choroidal malformations (n=87 WT, n=86 *rasa1a^+/-^* and n=36 WT, n=68 *ephb4a^+/-^*) **E.** Quantification of the diameter of the posterior segment in larvae with no choroidal malformation (n=87 WT, n=53 *rasa1a^+/-^*) in *rasa1a^+/-^*larvae and (n=36 WT, n=48 *ephb4a^+/-^)* in *ephb4a^+/-^* larvae at 3 dpf (Mann–Whitney U test). Scale bars, 300 µm for (B) and 50 µm for (C).

We decided to use the zebrafish model to investigate the causality between the mutations and the formation of neurovascular defects. Indeed, the transparency and rapid neurovascular development of the zebrafish larvae make them models of choice to investigate vascular morphogenesis and blood flow dynamics in real-time. First, we generated a *rasa1a* zebrafish mutant allele by CRISPR/cas9-mediated genome editing, targeting a sequence in exon 2 of *rasa1a* (rasa1aulb28). In the mutant allele, a frameshift of 7 bp, induces a premature stop codon, as observed in patients^3,14,15,27^ (Sup. Figure 1A). Homozygous *rasa1a^-/-^* can reach adulthood, although at frequencies lower than the one predicted by Mendelian genetics, exhibit a smaller size than heterozygous rasa1a^+/-^ or wildtype (W)T zebrafish and are not fertile (Sup. Figure 1B). Second, we obtained an *ephb4a* mutant line from ZIRC (*ephb4a*^hu3445^), where a single point mutation in exon 4 leads to an early stop codon (Sup. Figure 1A). We exclusively used heterozygous mutants (*rasa1a^+/-^*, *ephb4a*^+/-^) and wild-type (WT, *rasa1^+/+^*, *ephb4^+/+^)* zebrafish in our study, as the patients are also heterozygous. We did not observe cardiac edema in 3 days post fertilization (3 dpf) rasa1a^+/-^ and ephb4a^+/-^ larvae and observed less than 15% cardiac edema at 5 dpf (Figure 1B). We only observed a few heterozygous mutant larvae with minor intersegmental vessel (ISV) vascular growth defects at 5 dpf (Sup. Figure 2). Homozygous mutant larvae displayed more frequently cardiac edema and ISV growth defects than the heterozygous mutant larvae at 3 dpf and 5 dpf (Figure 1B, Sup. Figure 2). Therefore, heterozygous larvae display a more stable and uniform general phenotype than homozygous larvae and are fertile.

At 72 hpf (hours post fertilization, hpf), both *rasa1a^+/-^* and *ephb4a^+/-^* mutants displayed similar structural malformations of the dorsal longitudinal vein (DLV, Figure 1C). We observed identical malformations of the DLV in rasa1a and ephb4a morphans (Sup. Figure 3). The DLV belongs to the dorsal venous system, as the vein of Markowski^28^, the precursor of the vein of Galen, and sits above the diencephalon, mesencephalon, and metencephalon. We therefore propose, as others did^25^, that the DLV is a functional analog of the developing vein of Galen^29^. After analyzing larvae from both heterozygous mutant lines, we describe two different structural defects of the DLV at 72 hpf: on one form, we observed the presence of dissociated vessels on the anterior segment with, sometimes, dissociated vessels on the posterior segment (∼40% of all *rasa1a^+/-^* larvae and ∼30% of all *ephb4a^+/-^* larvae). On the remaining heterozygous larvae (*rasa1a^+/-^*and *ephb4a^+/-^*), non-affected by a choroidal malformation, we observed a significantly more dilated posterior segment of the DLV compared to WT larvae (Figure 1C). So, heterozygous mutant larvae either develop choroidal malformations, mural malformations, or no phenotype on the DLV, at 72 hpf. These two structural malformations are reminiscent of the two clinical forms of the disease described in human patients: the form with multiple vessels on both the anterior and posterior segments resembles a choroidal malformation with numerous fistulas. In contrast, the form with a dilated posterior segment resembles the mural malformation, with a dilation of the median vein.

Patients suffering from VGAMs often develop high-output heart failure^30^, a critical factor for the survival of neonate patients affected by choroidal malformations. We examined the cardiac function of *rasa1a^+/-^* and *ephb4a^+/-^* larvae at 72 hpf. We observed a significant increase in the stroke volume, corroborated by an increase in atrium and ventricle volumes during diastole (Figure 2A and B). We also observed increased blood flow velocity in the dorsal aorta, reflecting the increased heart output (Figure 2C). Therefore, both *rasa1a^+/-^*and *ephb4a^+/-^* mutants replicate the human disease’s genetics and major cardiovascular tissue defects with neurovascular malformations (fistulas or dilation) in the dorsal venous system as well as increased cardiac output.

**Figure 2:**
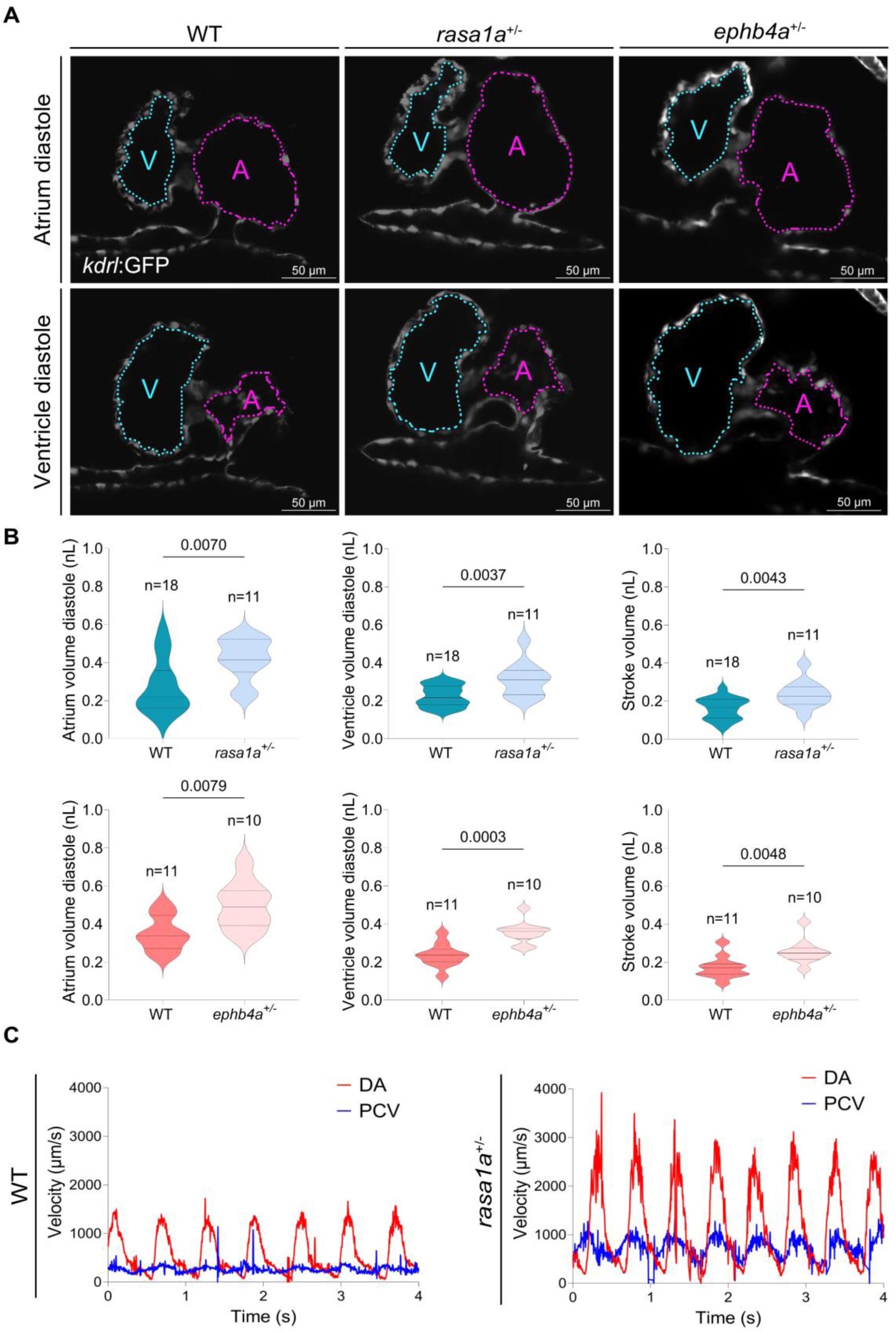
Characterization of the cardiac function of *rasa1a* and *ephb4a* mutants. **A.** Acquisition of *Tg(kdrl:GFP)* heart cycle of WT, *rasa1a*^+/-^ and *ephb4a^+/-^*larvae at 3 dpf. The atrium (A) is outlined in magenta and the ventricle (V) in cyan during the systole and the diastole. **B.** Quantification of the volume of atrium (A) diastole and ventricle (V) diastole and stroke volume (n=18 WT, n=11 *rasa1a*^+/-^ and n=11 WT, n=10 *ephb4a^+/-^*) at 3 dpf (Mann– Whitney U test). **C.** Velocity of red blood cells in the DA (red) and the PCV (blue) at 3 dpf for WT and *rasa1a^+/-^* larvae. Scale bars, 50 μm for (A).

### VGAMs result from a defective flow-mediated fusion and constriction of precursor vessels

The existence of multiple, perfused, and functional blood vessels in the malformation urged us to investigate its formation mechanism. We performed time-lapse imaging of the formation of the DLV in WT embryos. We observed a progressive fusion of two distinctly perfused precursor vessels within the first 4h following 46 hpf. Then, we observed the elongation and constriction of the fused vessel in the next 4h (Figure 3A, B, and C, supplemental movie 1). Endothelial cells from distinct precursor vessels interact by direct contact before the fusion of the two precursor vessels (supplemental movie 2). Initially, we observed the formation of a perfused donut-shaped structure. Cells then rearranged and migrated, fusing the donut and forming a collecting vessel with a single lumen; then, the blood vessel elongated and constricted to form the DLV itself.

**Figure 3:**
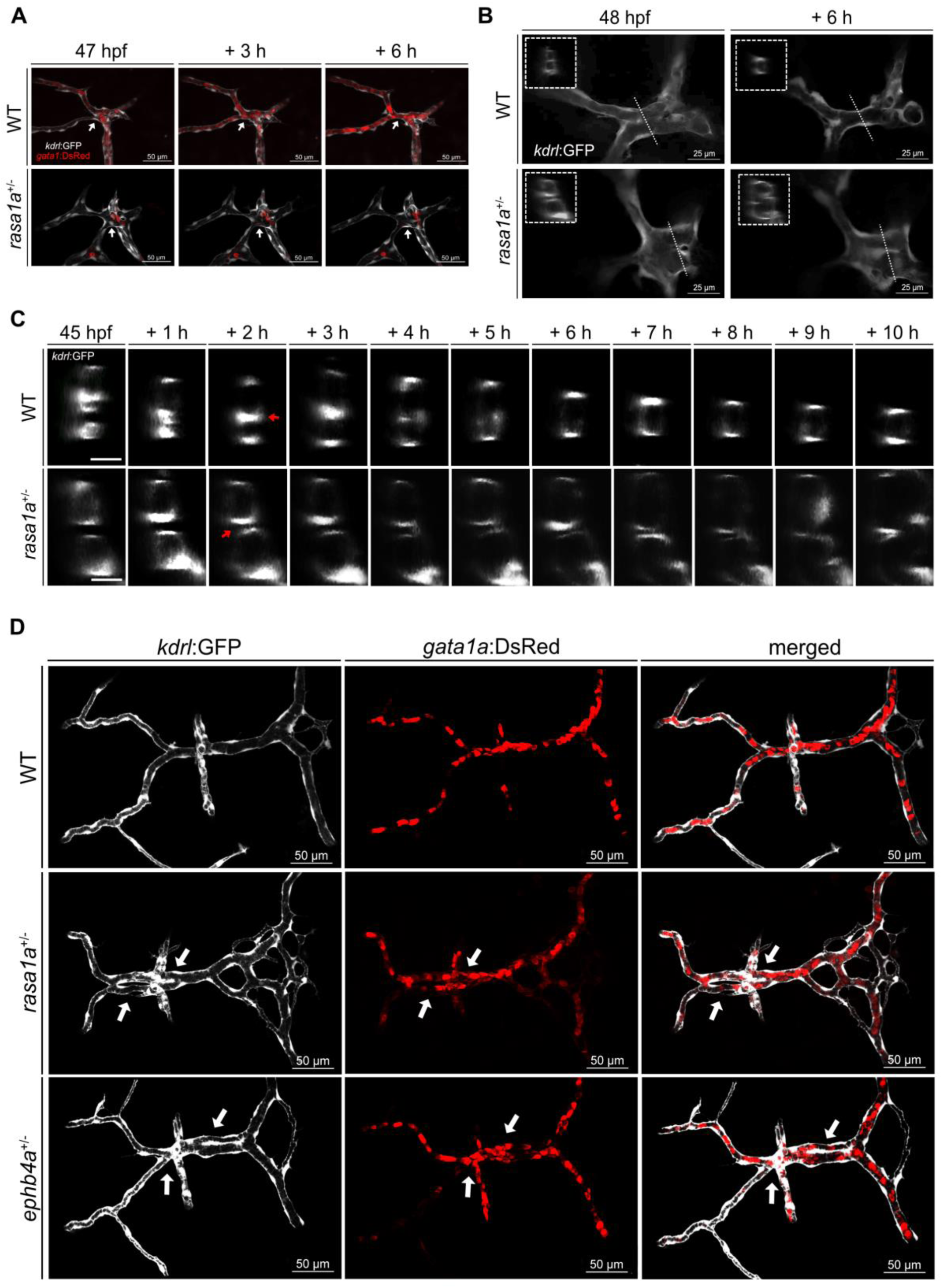
VGAMs are consecutive to a fusion and constriction failure during the development of the collecting vessel **A.** Maximum projection of the vessels and the red blood cells of WT and *rasa1a^+/-^*in *Tg(kdrl:GFP);(gata1a:DsRed)* double transgenics. The white arrows indicate the perfusion of two different streams to become one for WT larvae, while remaining as two separate streams in *rasa1a^+/-^* larvae. **B.** Observation of the formation of the anterior segment of the DLV at 48 and 54 hpf (collecting vessel, white square: transversal section at the location of the dashed line) **C.** Time lapse observation of the formation of anterior segment of the DLV (collector vessel in WT and *rasa1a^+/-^*), transversal section. The white arrows show the lumen total fusion for the WT and the partial one for the mutant. **D.** Maximal intensity projection of the dorsal view of the DLV of WT, *rasa1a^+/-^* or *ephb4a^+/-^* larvae with a choroidal malformation at 3 dpf. The white arrows indicate two perfused vessels on the anterior and 2 perfused vessels on the posterior segment of the DLV. Scale bars, 50 µm for (A) and (D), 25 µm for (B) and 10 µm for (C).

*rasa1a^+/-^* embryos display an ineffective vessel fusion process, developing distinct vessels (Figure 3A-C, supplemental movie 3). Indeed, the cells from the precursor vessels do not interact and fuse, as observed in the WT embryos. The distinction between the mural and choroidal malformations, described in Figure 1, most likely stems from which step of the DLV formation process (cell-cell interaction and fusion vs. constriction and elongation of the fused vessel) is ineffective. The absence of cell-cell contact, cell reorganization, and fusion of the initial donut-shaped structure leads to choroidal malformations with multiple fistulas in both the anterior and posterior portions of the dorsal venous system, as the precursor vessels that did not fuse further developed as perfused fistulas (Figure 3D). On the other hand, defective constriction of the fused vessels at later stages develops into a mural malformation. Both are distinct phenotypes. The fusion process requires that some endothelial cells, at the interface of two joining vessels (arrow, Figure 3C), perceive and integrate mechanical stimuli generated from the two pulsatile bloodstreams for the fusion to be successful, with blood flow mechanotransduction as a potentially important driver of the fusion process.

Therefore, we assessed whether blood flow contributed to DLV development in WT and mutant zebrafish. We observed that the preliminary vascular structure leading to the formation of the DLV was already perfused at 36 hpf (Sup. Figure 4A) and that the choroidal malformation consists of perfused fistulas (Figure 3F). This data shows that blood flow is present before DLV formation and might contribute as a mechanical stimulus to developing the DLV. We used two different approaches to gain insight into the role of flow in this specific vascular development program. First, we stopped blood flow by injecting a morpholino targeting *tnnt2a* at a low concentration (0.25 ng/μL) to stop heartbeat from the beginning of the cardiovascular development, with no blood flow within the vascular system of the developing embryo^31–33^. While most of the zebrafish embryo vascular system develops as it should from a structural point of view (Figure 4A), as flow is not entirely required for its primary development^34^, we observed a complete lack of development of the DLV (Figure 4B) of both WT and mutant embryos. Indeed, in embryos of both genotypes, the DLV remained entirely undeveloped with no fusion or elongation of the precursor vessels, indicating that both the formation of the normal DLV and the malformations in mutant embryos require blood flow to form. We confirmed this observation with a second experiment, where we incubated the embryos in 0.65 mg/ml of tricaine right before the beginning of the fusion process at 47 hpf, stopping blood circulation only right before this crucial step. As observed with *tnnt2a* morpholinos, blocking blood flow pharmacologically impaired the development of the DLV (Figure 4C).

**Figure 4:**
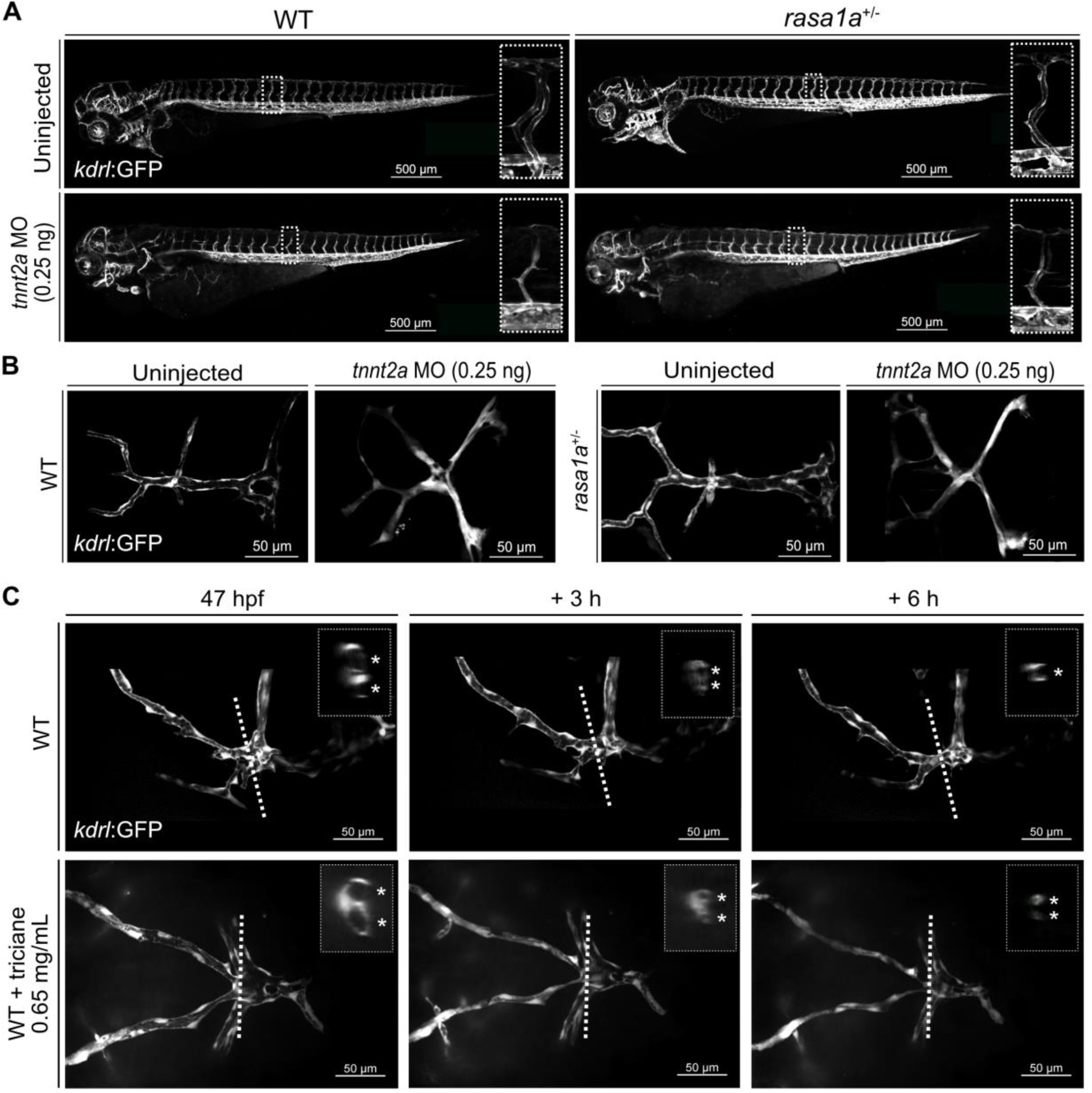
Rasa1 contributes to the formation of collecting vessels through a flow-mediated fusion of immature vessels **A.** Lateral views of WT and *rasa1a^+/-^* embryos injected with 0.25 ng *tnnt2a*-morpholino, at 3 dpf (white square: magnification of ISV) **B.** DLV of WT and *rasa1a^+/-^* embryos injected with 0.25 ng *tnnt2a*-morpholino, at 3 dpf. **C.** Stills of a time-lapse recording of WT DLV formation with and without tricaine overdose (0.65 mg/mL) (the white square is a transversal section at the location of the dashed line, and asterisks indicate the number of distinct vessels). Scale bars, 500 µm for (A) and 50 µm for (B) and (C).

We also counted the number of endothelial cells, as an increased cell number could be required to induce additional vessels in choroidal malformations. The analysis of the number of stained nuclei in *kdrl:nls.cherry* transgenic line did not reveal a significant difference in the number of endothelial cells in the DLV of WT compared to DLV with a choroidal malformation and multiple perfused vessels in *rasa1a^+/-^* mutants (Sup Figure 4B).

### The Dorsal Longitudinal Vein is a collecting vessel with singular hemodynamic properties

We observed stable, perfused fistulae in the DLV of *rasa1a^+/-^*and *ephb4a^+/-^* mutants, with no similar defects observed elsewhere in the systemic vasculature (Sup. Figure 2). We then decided to characterize the hemodynamic properties of this vascular bed more precisely and compare it to other systemic vessels. Tracking red blood cell velocities extrapolated hemodynamic parameters within zebrafish’s blood vessels. We observed a pulsatile flow with high shear stress in the dorsal aorta and a non-pulsatile flow with low shear stress in the posterior cardinal vein, as expected for an artery and a vein (Figure 5A). Interestingly, we measured a pulsatile flow with high shear stress in the DLV, almost similar to what is observed in the dorsal aorta (DA) and much higher to what is observed in the posterior cardinal vein (PCV) (Figure 5C). Frequency analysis of the pulsatility in DA determined a fundamental harmonic of 1.8 Hz, corresponding to the frequency of the beating heart. In comparison, the fundamental harmonic of the DLV was doubled at 3.8 Hz (Figure 5B). This surprising result led us to analyze the behavior of single red blood cells. We individually tracked red blood cells from each incoming vessel and labeled their tracks with a distinctive color for each inlet (green for the top and red for the bottom in the case of Figure 5D). It revealed independent tracks that were restrained to the side of each entry along the DLV as if they could not cross an imaginary border at the vessel’s center. This observation demonstrates the existence of streams of red blood cells within the DLV, which acts as a collecting channel where streams do not mix. As they do not mix, these streams possess their characteristics regarding pulsatility and, most likely, pressure and other hemodynamic parameters.

**Figure 5:**
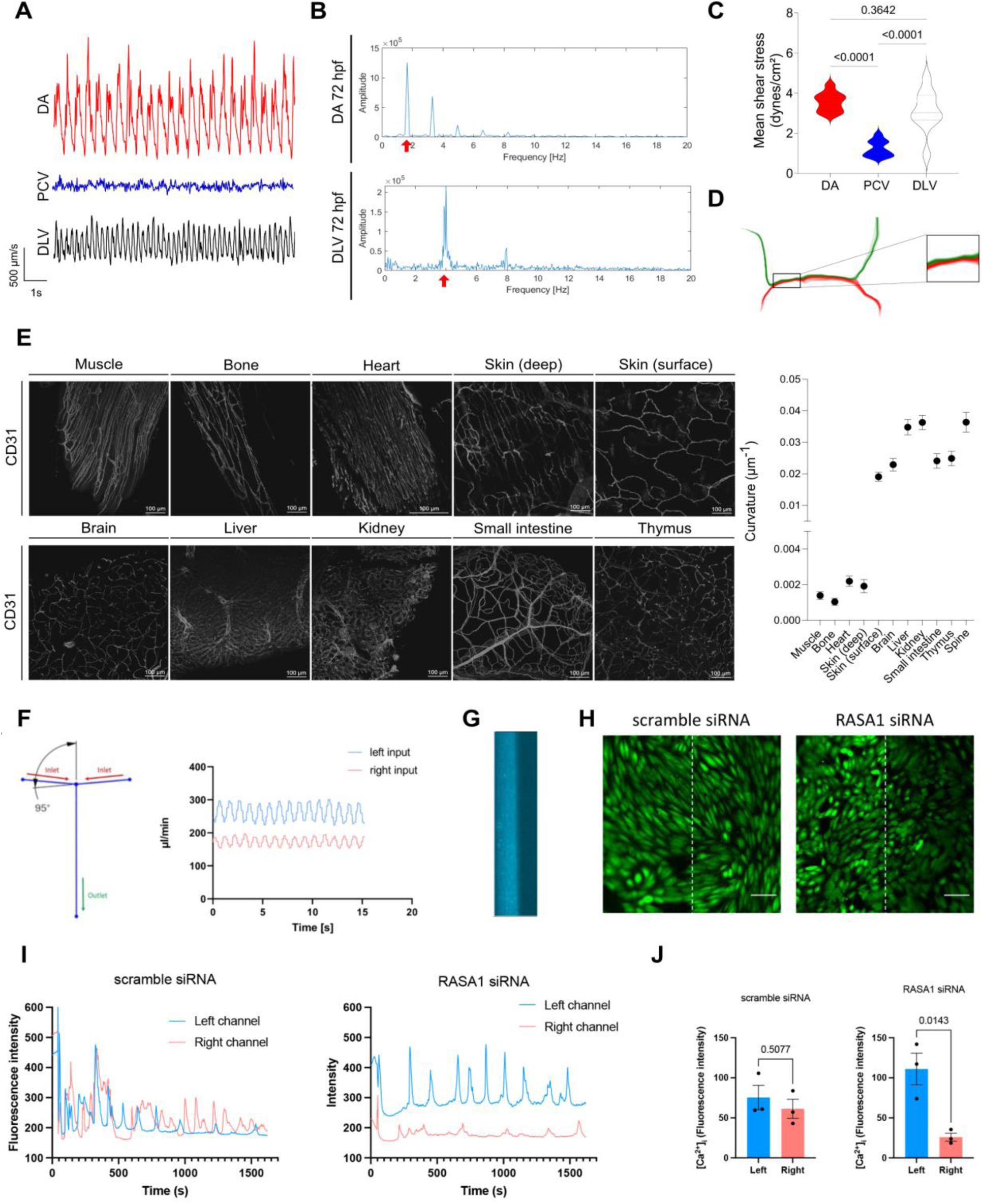
Analysis of blood flow and mechanosensation in a collecting channel **A.** Velocity of red blood cells in the dorsal aorta (DA), the posterior cardinal vein (PCV) and the dorsal longitudinal vein (DLV). **B.** Frequency analysis by Fourier transform of RBC velocity. **C.** Estimation of fluid shear stress intensities in each vessel. (n=10) **D.** Analysis of the trajectory of red blood cells originating from the top left entry (green) or the bottom left entry (red). **E.** Analysis of the curvature of capillaries in various organs transparized from mice with the iDISCO protocol and stained for CD31. The graph on the right displays the average curvature (mean +/- SEM) of 17 to 100 capillaries for each tissue. **F.** Description of the flow chamber and representative flow measures injected in each inlet. **G.** Capture of the channel within the flow chamber. The left inlet has been perfused with fluorescent blue beads, and the right inlet has non-fluorescent water. **H.** Representative images of intracellular calcium with a calcium reporter (Calbryte-520 AM) 25 mins after applying flow on HUVECs transfected with a scramble or a RASA1 siRNA. **I.** Intracellular calcium measurements in HUVECs transfected with a scramble or a RASA1 siRNA. One representative calcium trace is displayed over 30 mins from the collecting channel’s left or right side. **J.** Histograms of the average intensity of calcium oscillation (n=3, unpaired t-test). Scale bars, 100 µm for (E) and 50 µm for (H)

High-flow AVMs are frequently observed in two distinct congenital syndromes: HHT syndrome characterized by loss of function (LOF) genetic mutations affecting BMP signaling^35^; and CM-AVMs with LOF genetic mutations affecting *RASA1*^15^ or *EPHB4*^27^. Interestingly, HHT malformations mainly develop in specific vascular networks^36^ (brain, liver, lung, mucosae of the digestive system, and the superficial vascular network of the dermis). Similarly, CM-AVMs malformations have been described to develop in distinct vascular beds than HHT-related AVMs, namely in the skin’s deep vascular network, musculoskeletal vascular networks, and the developing vein of Galen^37^. We hypothesized that CM-AVMs develop in different vascular beds than AVMs of HHT patients, possibly due to a different structural organization and blood flow profiles within those respective vascular beds. We performed volumetric imaging of several mouse organs to compare the architecture of their vascular networks and identify possible common structural features among them. Strikingly, capillaries from muscles (cardiac or skeletal), bone, and the deep vascular network of the dermis, where CM-AVMs malformations preferentially develop in patients, shared a common structural feature: they consist of straight, elongated, collecting vessels with a similar architecture to the one observed in the DLV. Contrastingly, capillary networks of the brain, the superficial layer of the dermis, the kidney, the liver, the small intestine, and the thymus, where HHT-related malformations develop, displayed pronounced curvature of their capillaries (Figure 5E). The analysis of the curvature of these capillaries revealed two very distinctive groups, with more than an order of magnitude of difference for the curvature of the capillaries between HTT and CM-AVMs associated capillary beds. (Figure 5E). This difference suggests that the formation of those distinct capillary networks might stem from different angiogenesis programs and that hemodynamics would differ in each group to fulfill a different function. It is well known that curves in microscopic vessels generate centrifugal forces, which cause secondary flow vortices^38^, allowing the fluid to be mixed. In contrast, straight vessels similar to DLV might have different streams with poor mixing, a well-known issue in microfluidics^39^.

### Rasa1 contributes to the mechano-adaptation to non-homogeneous flow

Blood flow mechanotransduction involves numerous receptors and signaling pathways^1,13,40,41^. To investigate the contribution of Rasa1 to blood flow sensing, we developed a simplified in vitro PDMS model of the DLV, with a collecting channel perfused by two orthogonal inlets that can be perfused independently. We perfused the left inlet with a pulsatile flow of higher magnitude than the right intlet (Figure 5F), generating two fluid streams with distinct properties that do not mix inside the collecting channel, as observed in the DLV of the zebrafish larva (Figure 5G). Despite two streams with different mechanical parameters and different mechanical stimulation across the channel, WT cells transfected with a scramble siRNA exhibited a uniform calcium response in the left and right portions of the channel 25 minutes after the flow application, indicating that the endothelial cell monolayer responded uniformly to a non-homogenous mechanical stimulation, mitigating the differences (Figures 5H, 5I and 5J, supplemental movie 4). Consequently, endothelial cells can be subjected to different mechanical stimuli, which can be perceived and integrated to make the response uniform across a monolayer within a vessel through cell-cell communication. Strikingly, cells deficient for RASA1 exhibited a sharply different response on the left and right portions of the channel (Figures 5H, 5I, and 5J, supplemental movie 5). Calcium response is markedly and significantly higher in the left portion of the channel than in the right portion. Therefore, as a flow-sensitive organ, the endothelial monolayer is separated into two distinctive portions when Rasa1 expression is depleted. This indicates that Rasa1 contributes to flow-mediated cell-cell communication in an endothelial cell monolayer.

When organized in a monolayer, endothelial cells subjected to physiological laminar flow will align in the flow direction^42^, a unique endothelial cell feature associated with endothelial homeostasis^1,43^. We observed that HUVECs transfected with a siRNA against Rasa1 do not lose their ability to sense flow direction and to align in the flow direction, just as normal cells (Fig. 6A). Therefore, RASA1 specifically modulates only some aspects of the response to flow. Numerous signaling pathways are transduced in response to flow, including PI3K and ERK signaling. These two signaling cascades are known to be downstream of Rasa1 in response to chemical agonist stimulation, as Rasa1 is a RasGAP protein^44^. Therefore, we measured how Rasa1 knockdown would affect the flow-mediated activation of Akt and ERK. First, we measured how rasa1 knock-down affects VEGF-mediated activation of ERK and Akt, as VEGF is known to activate both pathways during angiogenesis^45^. As depicted in Figure 6B, we could not observe any significant difference between cells transfected with a scramble siRNA and cells transfected with Rasa1 siRNA in response to VEGF stimulation.

**Figure 6:**
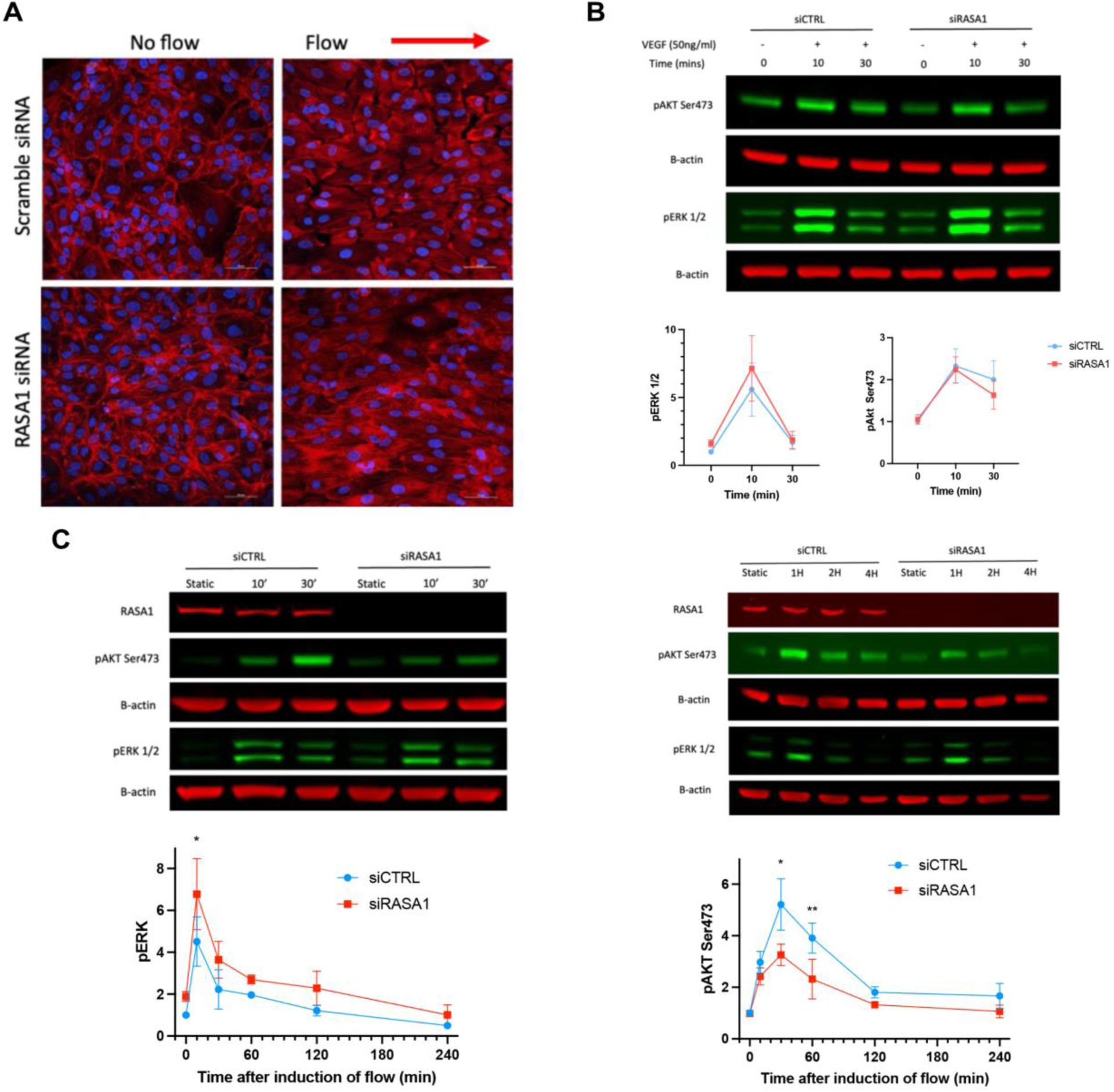
Identification of impaired flow mechanotransduction responses **A.** Endothelial cell monolayer organization after 16h of laminar flow (12 dynes/cm2) applied on HUVECS transfected with a scramble or RASA1 siRNA, the arrow indicates the direction of flow. Cell stained for f-actin with phalloidin (red) and the nuclei with DAPI (blue). **B.** Western blot of ERK1,2 (Thr202/Tyr 204) and Akt phosphorylation (Ser 473) in response to VEGF (50ng/mL) for 0, 10 and 30 mins (n=3, ANOVA, Sidaks multiple comparison post-test). **C.** Western blot of ERK1,2 (Thr202/Tyr 204) and Akt phosphorylation (Ser 473) in response to laminar flow (12 dynes/cm^2^) for 0, 10 and 30 mins, 1h, 2h and 4h (n=3, ANOVA, Sidaks multiple comparison post-test). Scale bar, 50 µm for (A).

On the other hand, Rasa1 knock-down cells exhibited defective flow-dependent activation of ERK, and Akt: ERK phosphorylation was slightly potentiated, while Akt phosphorylation by PI3K was significantly reduced after flow (Figure 6C). This contrasts with cells transfected with a scramble siRNA, where Akt and ERK are activated after applying flow, although with different kinetics. This observation confirms the specific involvement of Rasa1 signaling in flow mechanotransduction. It provides ground for attempting to correct pharmacologically the inadequate response to flow and, potentially, the defective flow-mediated fusion of immature vessels.

### Targeting the defective flow-mediated signaling reverses choroidal and mural malformations by restoring flow-mediated vessel fusion

We hypothesized that correcting the imbalance in flow activation of PI3K/ERK signaling would restore proper vessel fusion and constriction in the mutants. We treated WT, *rasa1a^+/-^* or *ephb4a^+/-^*zebrafish with either DMSO (used as a solvent), selumetinib (MEK1 inhibitor^46^, targeting ERK activation, 1 µM), YS-49 (PI3K activator^47–49^, restoring Akt activation, 1 µM), and a combination of selumetinib and YS-49 (1 µM each). We deployed two protocols to assess whether pharmacological treatment would restore DLV structure and mitigate the development of the malformation. The first protocol (Figure 7A) consisted in incubating the drugs from 36 hpf to 60 hpf, during the development of the DLV. We measured a significant reduction of the diameter of the DLV, with an increasing effect from selumetinib, YS-49, and then the combination of both drugs until returning to the values measured in WT zebrafishes (Figure 7B). We observed a reduction in the proportion of *rasa1a^+/-^* larvae displaying choroidal malformations (40% with DMSO) after using either selumetinib (25%) or YS-49 (20%) (Figure 7C). Combining both drugs did not improve the outcome compared to using a single drug. In *ephb4a^+/-^* larvae, we also observed similar results for the reduction of DLV diameter but the drugs were less efficient in reducing the number of larvae with choroidal malformations (Figures 7B and 7C). This experiment validates the hypothesis that a defective flow-mediated activation of MAPK and PI3K signaling contributes to malformation development.

**Figure 7:**
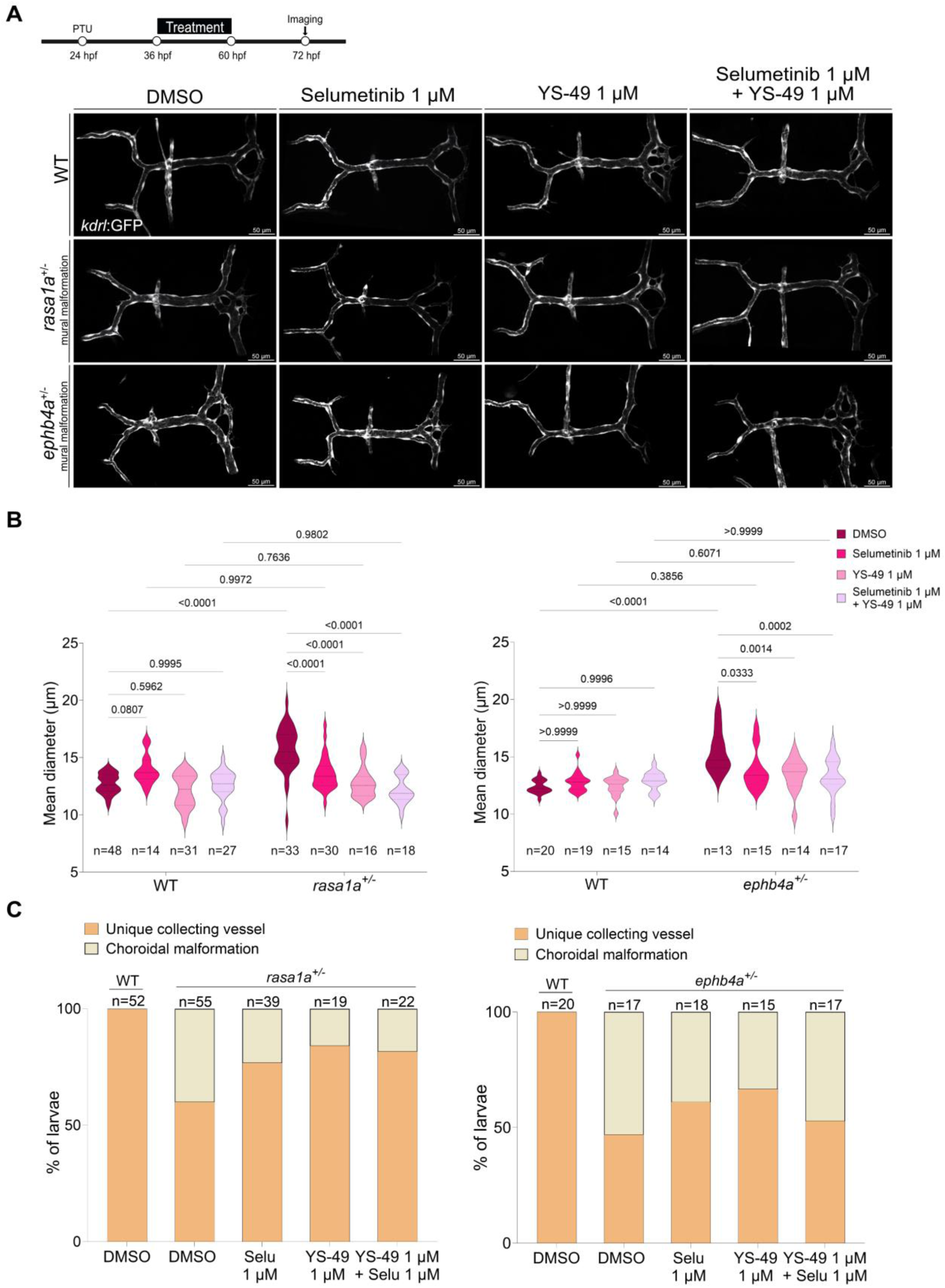
Pharmacological modulation of the PI3K or MAPK pathways prevents DLV malformations in *rasa1a^+/-^*and *ephb4a^+/-^.* **A.** Effect of the inhibition of ERK signaling (Selumetinib 1 µM) or activation of PI3K (YS-49 1 µM) or a combination of both during the development of the DLV (36 hpf until 60 hpf). Maximum projection of the DLV of *Tg(kdrl:GFP)* WT, *rasa1a^+/-^*and *ephb4a^+/-^* DLV with a mural malformation before and after the pharmacological treatment. **B.** Quantification of the average diameter of mural malformation after the pharmacology treatment of the WT, *rasa1a^+/-^* and *ephb4a^+/-^* larvae (left graphe : WT DMSO : n=48, *rasa1a^+/-^* DMSO: n=33, WT Selumetinib : n=14, *rasa1a^+/-^* Selumetinib: n=30, WT YS-49 : n=31, *rasa1a^+/-^* YS-49 : n=16, WT Selumetinib + YS-49 : n=27 and *rasa1a^+/-^* Selumetinib + YS-49 : n=18 / right graph : WT DMSO : n=20, *ephb4a^+/-^* DMSO: n=13, WT Selumetinib : n=19, *ephb4a^+/-^* Selumetinib: n=15, WT YS-49 : n=15, *ephb4a^+/-^* YS-49 : n=14, WT Selumetinib + YS-49: n=14 and *ephb4a^+/-^* Selumetinib + YS-49 : n=17). **C.** Quantification of the percentage of choroidal malformation after the pharmacology treatment of the WT, *rasa1a^+/-^* and *ephb4a^+/-^* larvae (left graph : WT DMSO : n=52, *rasa1a^+/-^* DMSO n=55, *rasa1a^+/-^*Selumetinib : n=39, *rasa1a^+/-^* YS-49 : n=19, *rasa1a^+/-^* Selumetinib + YS-49 : n=22 / right graph : WT DMSO : n=20, *ephb4a^+/-^* DMSO n=17, *ephb4a^+/-^* Selumetinib : n=18, *ephb4a^+/-^* YS-49 : n=15, *ephb4a^+/-^* Selumetinib + YS-49 : n=17). Statistical analysis: Two-way ANOVA with multiple comparisons. Scale bars, 50 µm for (A).

We then tested whether the drugs might reverse the two malformations as a potential therapeutic approach. To test this, we treated larvae with the different drugs at 72 hpf for 24h, when the malformation was fully formed, and monitored the drug’s effect on the mural or choroidal malformations after 24h (Figure 8A).

**Figure 8:**
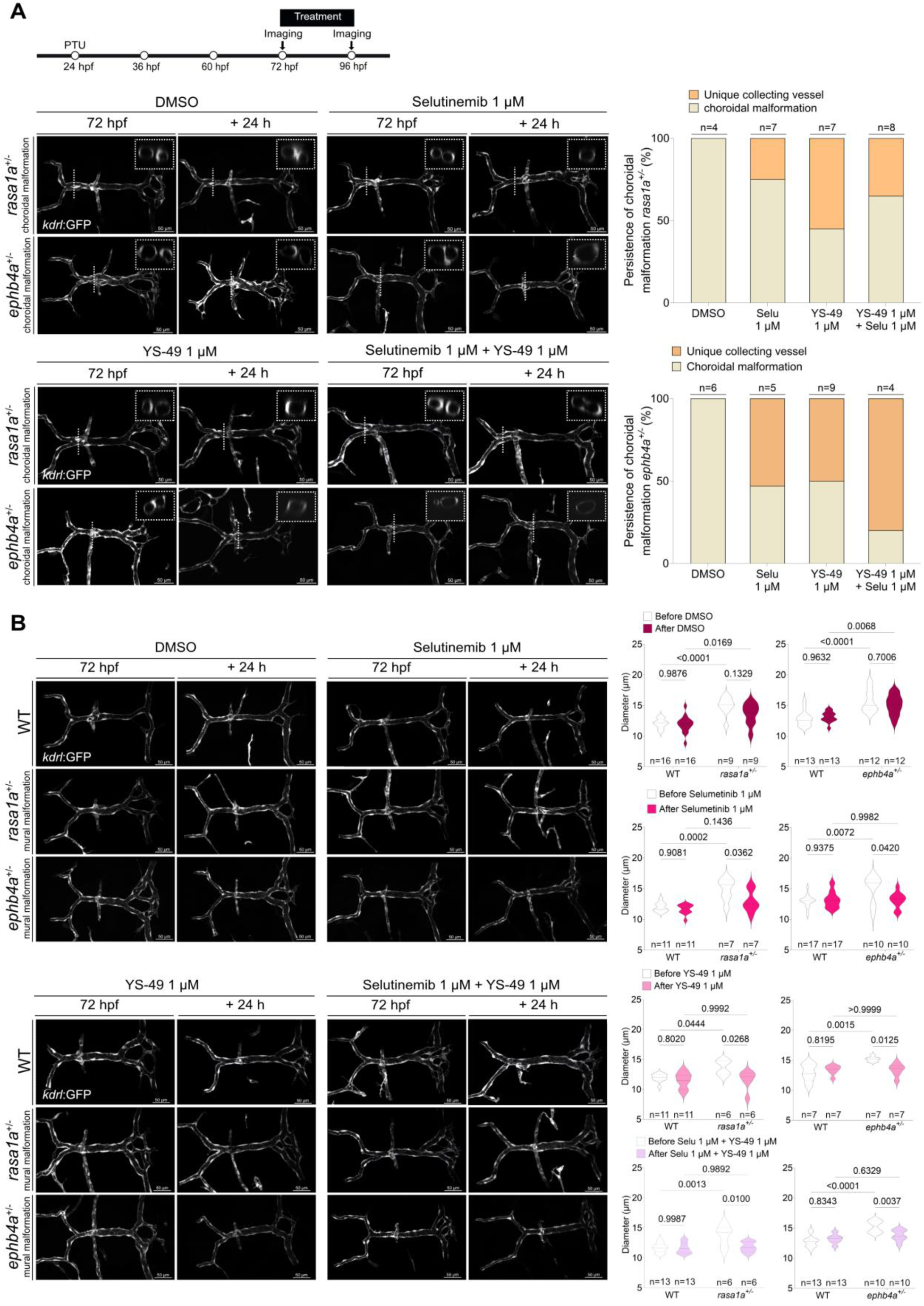
Targeting PI3K or MAPK pathways reverses DLV malformations **A.** Effect of the inhibition of ERK signaling (Selumetinib 1 µM) or activation of PI3K (YS-49 1 µM) or a combination of both after the development of the DLV or the malformations (72 hpf until 96 hpf). Maximum projection of the DLV from *Tg(kdrl:GFP)* WT, *rasa1a^+/-^*and *ephb4a^+/-^* with a choroidal malformation before and after the pharmacological treatment on the same larvae (72 hpf and 96 hpf). Within the small white square is a transversal section at the location of the dashed line showing the vessels structure. Quantification of the percentage of choroidal malformation before and after the treatment of *rasa1a^+/-^* larvae (DMSO : n=4, Selumetinib : n=7, YS-49 : n=7 and Selumetinib + YS-49 : n=8) and *ephb4a^+/-^* (DMSO : n=6, Selumetinib : n=5, YS-49 : n=9 and Selumetinib + YS-49 : n=4). **B.** Effect of the inhibition of ERK signaling (Selumetinib 1μM), activation of PI3K (YS-49 1μM) or a combination of both (YS-49 and Selumetinib) after the development of the DLV or the malformations (72 hpf until 96 hpf). Maximum projection of the DLV from *Tg(kdrl:GFP)* WT, *rasa1a^+/-^* and *ephb4a^+/-^* with a mural malformation before and after the pharmacological treatment on the same larvae (72 hpf and 96 hpf). Quantification of the average diameter of the mural malformation before and after the treatment of the WT, *rasa1a^+/-^* (DMSO : n=16 WT, n=9 *rasa1a^+/-^*; Selumetinib : n=11 WT, n=7 *rasa1a^+/-^*, YS-49 : n=11 WT, n=6 *rasa1a^+/-^* and Selumetinib + YS-49 : n=13 WT, n=6 *rasa1a^+/-^*) and *ephb4a^+/-^*larvae (DMSO : n=13 WT, n=12 *ephb4a^+/-^*; Selumetinib : n=17 WT, n=10 *ephb4a^+/-^;* YS-49 : n=7 WT, n=7 *ephb4a^+/-^*and Selumetinib + YS-49 : n=13 WT, n=10 *ephb4a^+/-^*). Statistical analysis: ANOVA with Tukey’s test for multiple comparisons. Scale bars, 50 µm for (A) and (B).

Interestingly, choroidal malformations remained stable from 72 hpf to 96 hpf in the absence of treatment (DMSO), highlighting the stability of these fistulae. The administration of either selumetinib, YS-49, or the combination of both reversed the choroidal malformation in nearly 50% of the treated *rasa1a^+/-^* larva, with a more marked effect of YS-49 (Figure 8B). The impact of the drugs was more pronounced on *ephb4a^+/-^* larvae, with more than 50% of larvae restoring vessel fusion with either selumetinib or YS-49 and 75% of larvae with a combination of both molecules (Figure 8B). It is, therefore, possible to trigger the fusion of established choroidal malformations.

The drugs’ impact on enlarged mural malformations in *rasa1a^+/-^* or *ephb4a^+/-^*larvae was also positive when using selumetinib, YS-49, or the combination of both molecules (Figure 8C), as we could measure a significant reduction of the diameter of the enlarged DLV. This suggests that the constrictive process could also be restored in enlarged mural malformations.

## Discussion

Vascular remodeling is an essential feature of vascular embryogenesis and homeostasis. Flow mechanosensing is one of the main drivers of lumen formation^50^, vascular remodeling, and vessel homeostasis^8^, yet, the specific nature of mechanosensing programs still needs to be better understood and mapped out. We know that the nature of the flow within a developing vessel will determine the nature of the remodeling program^1,13,40^. Pruning of lowly or non-perfused, immature capillaries redirects endothelial cells toward perfused areas^5,51,52^. On the other hand, forming pillars within a developing lumen, a process called intussusception, requires low perfusion to stabilize the pillar^53^. Then, it has been observed that highly perfused capillaries could undergo reverse intussusception or, simply put, fuse^54^. This later process, the flow-mediated fusion of immature vessels, has been directly linked to VE-cadherin activation^55^, a cell-cell transmembrane receptor contributing to a major flow mechanosensory complex^56,57^ and endothelial cell homeostasis. Still, this mechanism cannot explain the specific nature of the flow-mediated fusion of immature vessels, as VE-Cadherin phosphorylation is also observed in other flow-dependent remodeling programs^58,59^ and because VE-cadherin phosphorylation directly affects endothelial junctions formation and, therefore, the formation of the junctional mechanosensor, a *primum movens* in flow mechanotransduction.

Working with an animal model that is RASA1 or EPHB4 deficient has proven difficult, as seen in the embryonic lethality of the knock^60,61^￼. The two zebrafish mutants used in this study recapitulate the phenotype observed in *ephb4* ^25^, with the generation of fistulae in the dorsal venous system of the cerebral vasculature. We reproduced these results by using morpholinos targeting *rasa1a* or *ephb4a*. Another group has generated multiple *rasa1a* and *rasa1b*^62^ mutant alleles but did not describe the DLV morphology. Yet, they described malformations of the caudal venous system at an earlier stage of the larval development in double *rasa1a/rasa1b* mutants.

Our study provides further evidence for vessel fusion as the necessary process to form collecting vessels that drain multiple high-flow feeding vessels. The existence of different mechanical stimuli during the fusion process necessitates an integration and adaptation of the endothelial cells, which would have to act as a single tissue despite the differences across the monolayer. In the absence of Rasa1, cells cannot mitigate multiple mechanical signals from blood flow, reinforcing the stabilization of multiple single fistulas rather than the formation of a single collecting vessel by fusion. Failure to fuse could also be explained by a defective flow-dependent cell-cell communication, maybe through EphB4, the other mutated gene in VGAMs^27^ , or a defective flow-mediated signaling cascade. Indeed, calcium waves cannot propagate anymore across the endothelial monolayer. The exact molecular culprit remains to be identified. Others have proposed a defective collagen IV secretion, which might contribute to stabilizing the fused vessel^63^.

We also describe specific vascular beds where such collecting vessels are present, namely, musculoskeletal capillaries, deep dermal capillaries, the vein of Markovski, and myocardial capillaries. The developmental program of these vessels most likely involves flow-dependent vessel fusion, as we could not observe any development of the DLV in the absence of perfusion of the developing vascular network and because those vessels share structural features with the DLV and are frequently associated with lesions in CM-AVM syndrome. Many vascular diseases have been linked to specific flow sensing mechanisms: atherosclerosis develops in areas of multi-directional flow, and from a defective perception of flow direction^42,64,65^, HHT-associated AVMs result from a defective flow-mediated stabilization of vessel development^9,66,67^ , inward remodeling, and hypertensive diseases to low-flow sensing^12,68^. Our work provides the first evidence for a pathological consequence of a defective flow-mediated fusion angiogenic program. Therefore, it identifies Rasa1 and EphB4 as important molecular actors of this poorly understood physiological mechanism in angiogenesis. Interestingly, another group has reported vascular anomalies in the adult myocardial vascular network of the mouse but not the skeletal muscle after an endothelial-specific deletion of EphB4^69^.

Flow properties vary across the vascular system and depend highly on the local architecture. At the microvascular level, fluid flow is affected by architectural factors such as curvature^38^ and the nature of blood as a fluid^70^, and, therefore, we might postulate that physics within capillaries is different from physics in larger vessels. Further studies must be conducted to appreciate the exact nature of biomechanics within capillaries fully. We identified two subgroups of microvascular networks differentiated by their curvature, which correlate with the localization of lesions either in CM-AVMs or HHT syndromes, indicating that flow properties and geometrical characteristics of the affected vascular network are probable contributors to the location of a lesion. Further, it demonstrates that mechanotransduction pathways are differentially activated and impacted based on the different hemodynamics or genetic anomalies, explaining the differences across vascular remodeling defects in disease.

Clinically, VGAMs are occasionally detected on antenatal ultrasound scans (from about 25 weeks of gestation). More commonly, VGAMs are diagnosed at birth. Often, delivery and the first 24 hours are unremarkable. More significant shunts may rapidly deteriorate with progressively congestive heart failure (CHF), leading to multiorgan failure. Occasionally, children present later in childhood with macrocrania or prominent facial veins secondary to venous outflow obstruction. Embolization is the first line of treatment for VGAMs and is usually performed in several sessions with glue (liquid embolic agent). A good outcome is anticipated when this treatment is performed before significant brain damage has occurred. In a recent meta-analysis^26^, the result was favorable in 68% of patients and unfavorable in 31%, with a mortality rate of 16%. The worst prognosis was seen, as expected, in babies with the largest shunts, presenting as neonates with severe high-output heart failure. Despite improvements in endovascular technologies and neonatology, these results compare unfavorably with those obtained in older children and adults with arteriovenous shunts. Moreover, a recent study has shown that long-term outcomes are less favorable than short- and mid-term outcomes, even without encephalomalacia at birth^26^. These patients will develop neuropsychological disorders and neurodevelopmental alterations. Therefore, current VGAMs management has significant limitations in patients with a severe clinical presentation, and there is a need for potential pharmacological support to decrease the potential brain parenchymal destruction by venous hyper-pressure, decrease congestive heart failure, increase the success rate of endovascular procedures and reduce the late neurodevelopmental alterations. We identified PI3K activators and MAPK inhibitors as potential pharmacological modulators of VAGM formation. Interestingly, PI3K activators, in particular, could reverse an already stably formed choroidal lesion, providing the first preclinical evidence for a combinatory therapeutical approach in managing VGAMs with a poor prognostic. Stabilizing neonates pharmacologically should now be considered, and it could positively support their development until they are strong enough to undergo an endovascular procedure.

Improving the care of these neonates required a deeper understanding of the etiology, which needed the contribution of various disciplines. Indeed, by identifying the precise nature of the defective tissular mechanism leading to the malformations in a zebrafish model and, for the first time, identifying specific molecular actors of flow-mediated vessel fusion, we enable the rapid development of targeted strategies not only for VGAMs but also for other types of diseases triggered by vascular remodeling.

## Figures legends

**Supplemental Figure 1:**
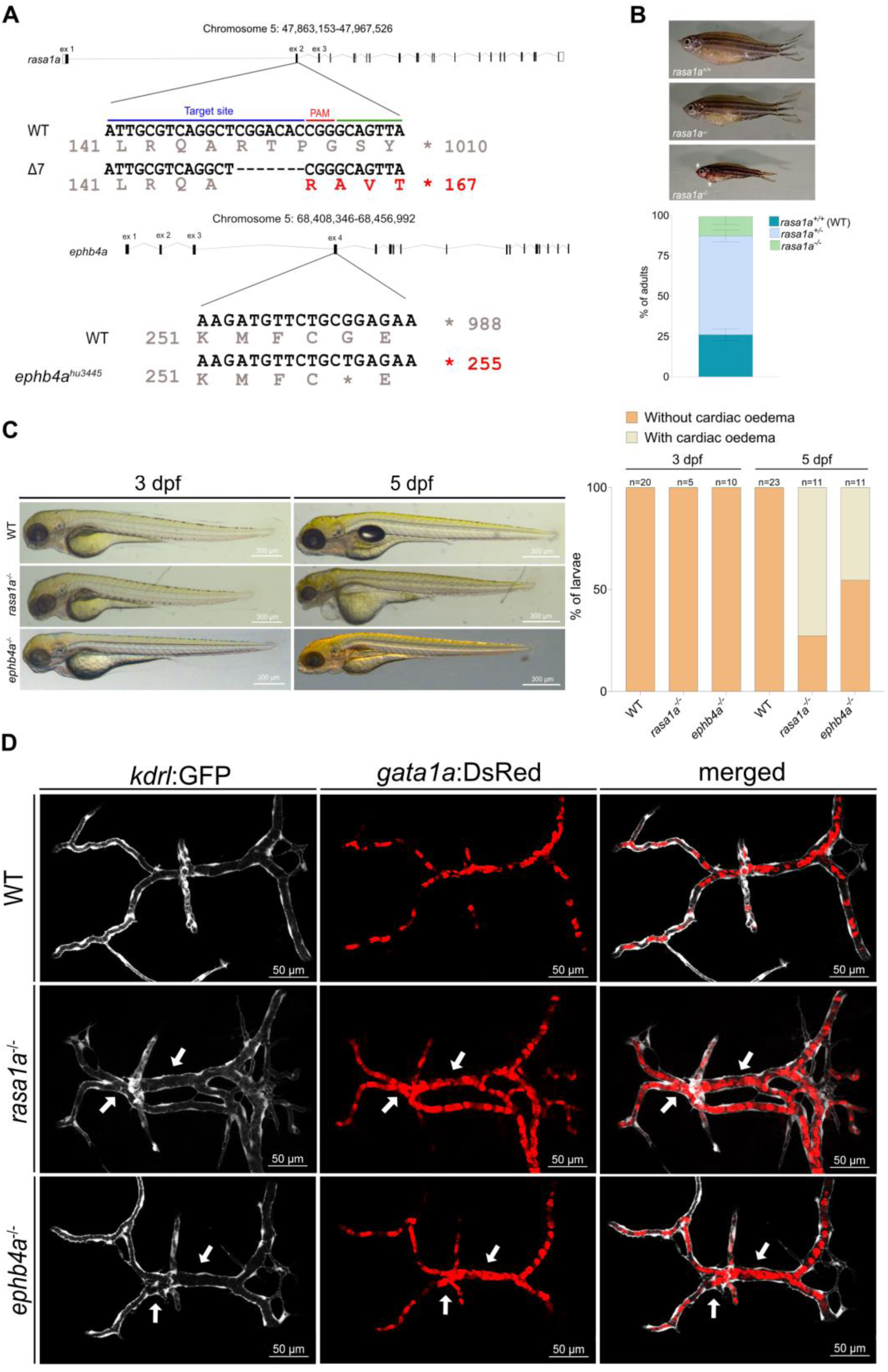
A. Characterization of the *rasa1a^-^* and *ephb4a* mutant alleles. PAM, protospacer adjacent motif. For *rasa1a*, The CRISPR/Cas9 target site is located in exon 2 of *rasa1a*. The mutant allele harbors a deletion of 7 nucleotides resulting in an early stop codon.. For *ephb4a*, a single point mutation is located in exon 4 of *ephb4a* and induces a stop codon. **B.** Lateral views of the adult WT and *rasa1a* mutants (asterisk on hemorrhagic spots). Genotyping of adult zebrafish derived from 4 different *rasa1a^+/-^* incrosses (mean +/- SEM) **C.** Lateral views of WT, *rasa1a^-/-^* and *ephb4a^-/-^* larvae at 3 dpf or 5 dpf. Quantification of cardiac edema in WT, *rasa1a^-/-^* and *ephb4a^-/-^* larvae at 3 dpf (n=20 WT, n=5 *rasa1a^-/-^* and n=10 *ephb4a^-/-^*) and 5 dpf (n=23 WT, n=11 *rasa1a^-/-^* and n=11 *ephb4a^-/-^*). **D.** Maximal intensity projection of the dorsal view of Tg(*kdrl:GFP);(gata1a:DsRed*) of the DLV of WT, *rasa1a^-/-^* and *ephb4a^-/-^* larvae with a choroidal malformation at 3 dpf. The white arrows indicate two perfused vessels on the anterior and/or 2 perfused vessels on the posterior segment of the DLV. Scale bars, 300 µm for (C) and 50 µm for (D).

**Supplemental Figure 2:**
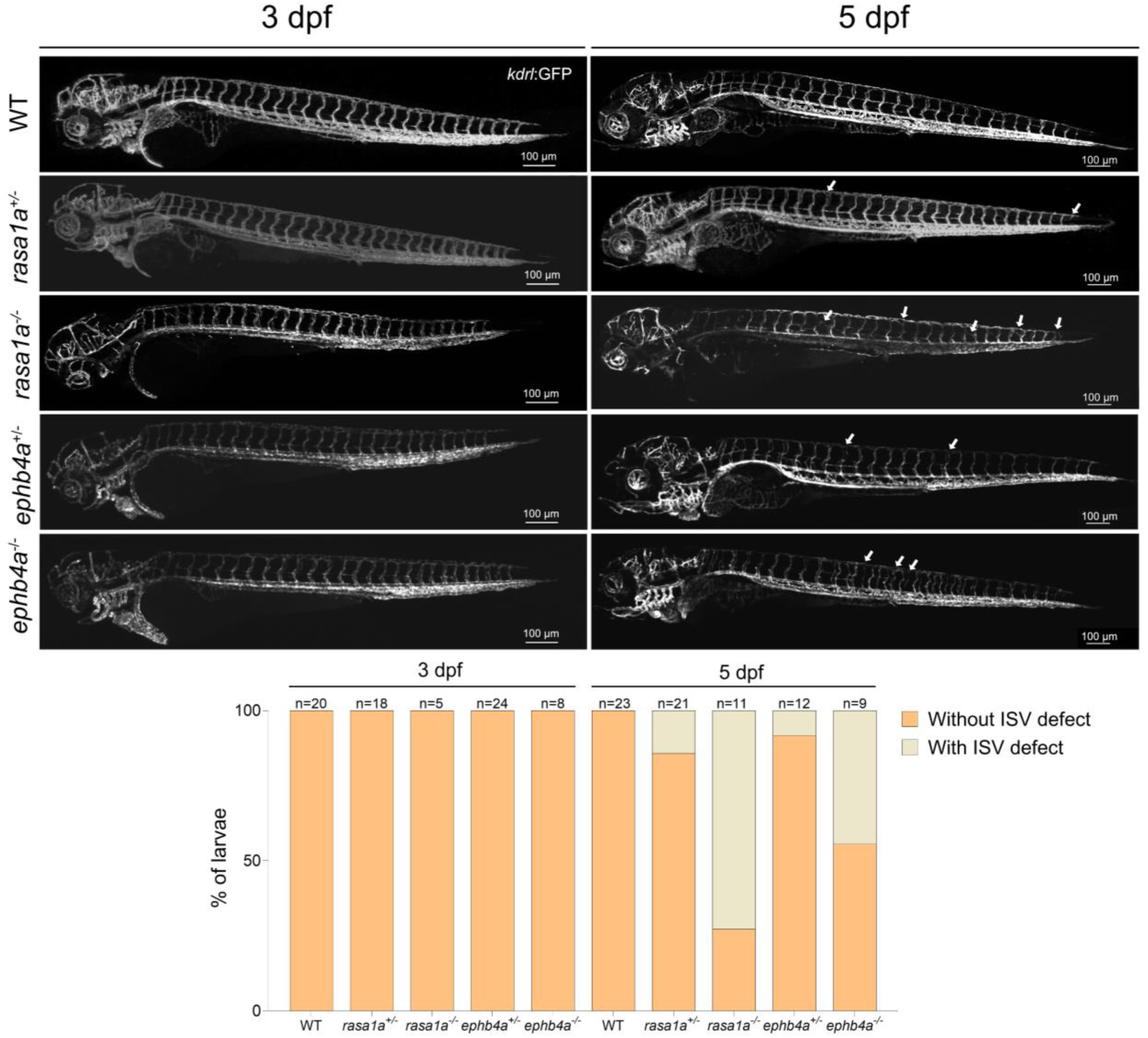
Maximal intensity projection of a z-stack of WT, *rasa1a^+/-^*, *rasa1a^-/-^*, *ephb4a^+/-^* and *ephb4a^-/-^* Tg(*kdrl*:GFP) larvae at 3 dpf (n=20 WT, n=18 *rasa1a^+/-^* n=5 *rasa1^-/-^*, n=24 *ephb4a^+/-^* and n=8 *ephb4a^-/-^*) or 5 dpf (n=23 WT, n=21 *rasa1a^+/-^* n=11 *rasa1^-/-^*, n=12 *ephb4a^+/-^* and n=9 *ephb4a^-/-^*). White arrows indicate regions of ISV defects, such as duplication or lack of perfusion. Quantification of ISV malformations in WT, *rasa1a^+/-^*, *rasa1a^-/-^*, *ephb4a^+/-^* and *ephb4a^-/-^* larvae at 3 dpf and 5 dpf. Scale bars, 100 µm.

**Supplemental Figure 3:**
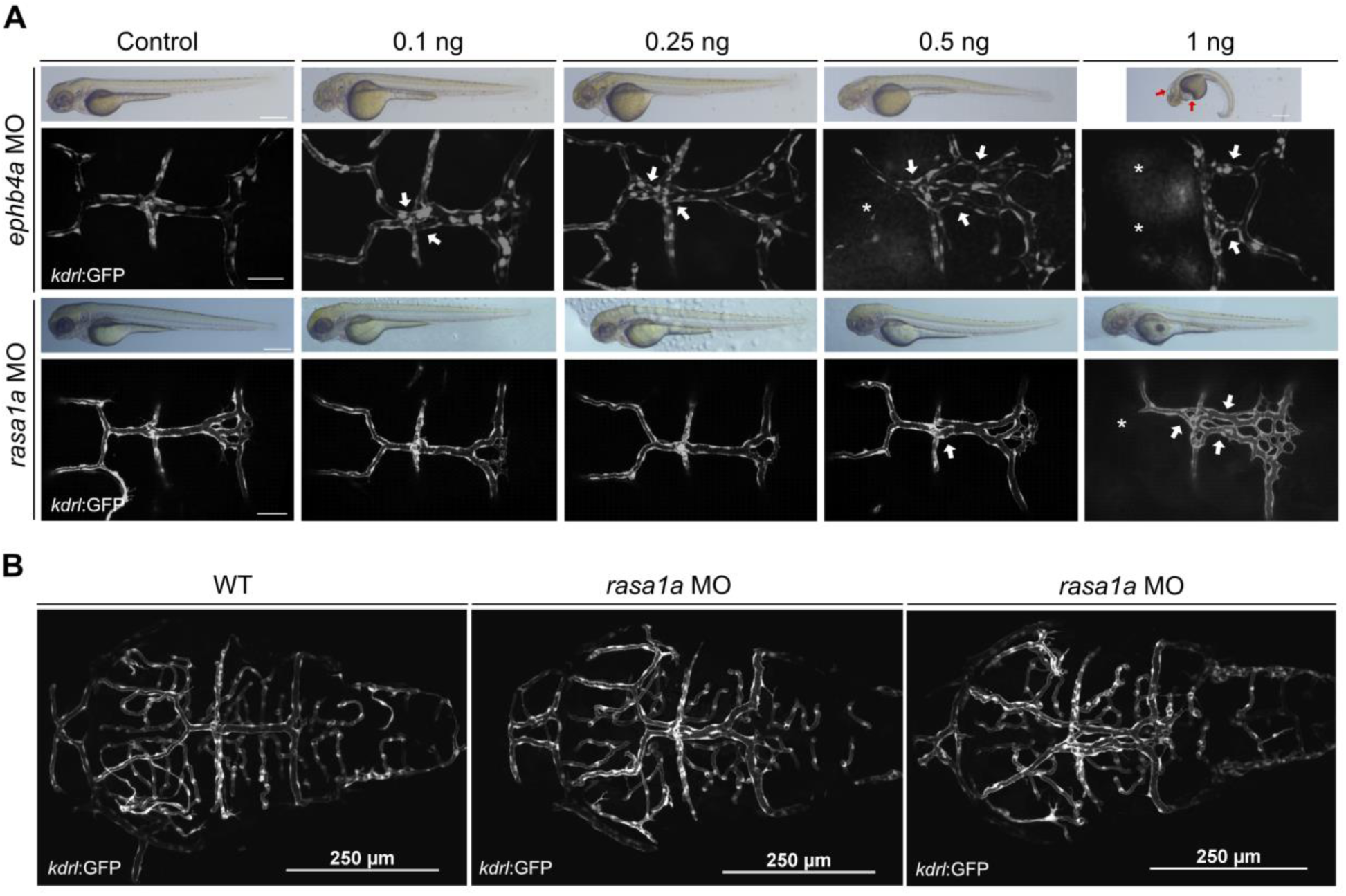
Lateral views of larvae at 3 dpf and maximal intensity projection of *Tg(kdrl:GFP)* of DLV in WT larvae at 3 dpf injected with different doses of *ephb4a* and *rasa1a* morpholinos. Injection of *ephb4a* morpholino at 0.5 ng and *rasa1a* morpholino at 1 ng resulted in duplication or triplication of the DLV (white arrow) and absence of the mesencephalic veins (MsV) (asterisk). At higher concentrations of *ephb4a* morpholino, cardiac and cerebral edema (red arrow) were observed. **B.** Dorsal view of the brain vasculature network at 3 dpf of WT embryos injected with 1 ng of morpholino against *rasa1a*. Scale bars, 300 µm and 50 µm for (A) and 250 µm for (B)

**Supplemental Figure 4:**
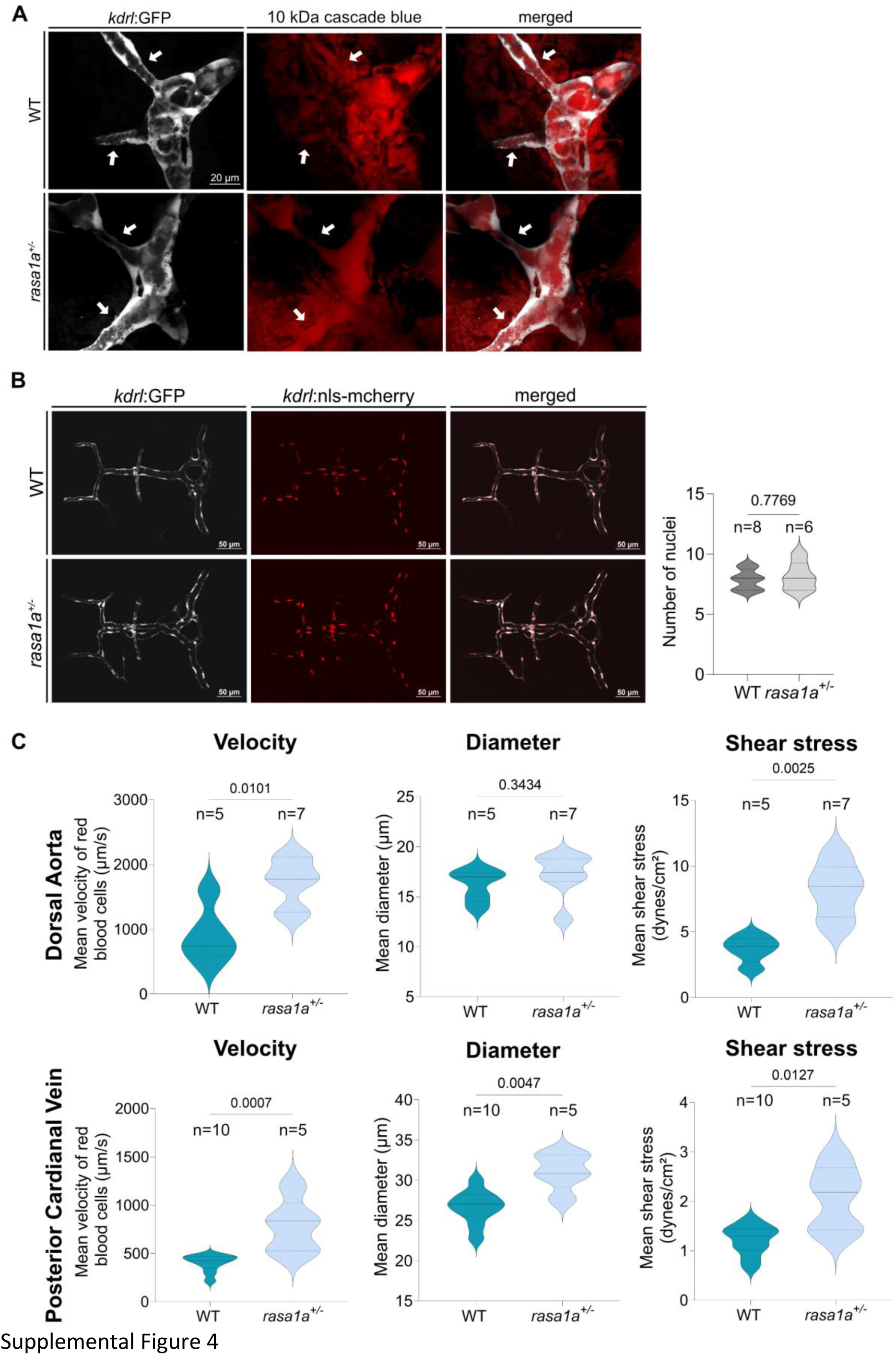
Analysis of vessel perfusion by fluorescent tracer injection (10kDa cascade blue) into the duct of Cuvier at 36 hpf. The preliminary vascular structure leading to the formation of the DLV was perfused at 36 hpf in WT and *rasa1a^+/-^* embryos. **B.** Maximum projection of the vascular network of the DLV in WT and *rasa1a^+/-^ Tg(kdrl:nls-mCherry);(kdrl:GFP)* double transgenics. Quantification of the number of nuclei in the DLV (n=8 WT, n=6 *rasa1a^+/-^*, Mann–Whitney U test). **C.** Quantification of DA and PCV blood flow parameters : velocity (DA : n=5 WT, n=7 *rasa1a^+/-^*; PCV : n=10 WT, n=5 *rasa1a^+/-^*), diameter (DA : n=5 WT, n=7 *rasa1a^+/-^* ; PCV : n=10 WT, n=5 *rasa1a^+/-^*) and shear stress (DA : n=5 WT, n=7 *rasa1a^+/-^* ; PCV : n=10 WT, n=5 *rasa1a^+/-^*) in WT and *rasa1a^+/-^* at 3 dpf (Two-way ANOVA with multiple comparisons). Scale bars, 20 µm for (A) and 50 µm for (B)

**Supplemental movie 1 and 2:** Timelapse imaging of transversal sections of *Tg(kdrl:GFP)* to observe DLV fusion in WT (movie 1) and *rasa1a^+/-^* larvae (movie 2).

**Supplemental movie 3 and 4:** Timelapse imaging of the DLV in WT *Tg(kdrl:ras-mCherry)* to observe endothelial cell migration without nuclear visualization (movie 3) or in WT *Tg(kdrl:GFP);(kdrl:ras-mCherry)* double transgenics for nuclear visualization (movie 4) during the fusion process.

**Supplemental movies 4 and 5:** Intracellular calcium changes on a monolayer of control HUVECs (movie 4) and HUVECs transfected with a rasa1 siRNA (movie 5) after application of flow in a chamber perfused by two inlets.

**Supplemental movies 6:** Timelapse imaging of *Tg(kdrl:GFP)* to observe the DLV formation in WT larvae.

## Acknowledgments

The authors thank Sabine Costagliola and Valérie Wittamer (IRIBHM, Université libre de Bruxelles) for their guidance throughout the project and for using their zebrafish husbandry. The authors would like to thank David Rodriguez for his help in analyzing the properties of the blood flow in zebrafish.

## Materials and methods

### Clinical imaging of VGAMs

Initial imaging diagnosis of VGAM are usually made during the third trimester with color Doppler ultrasonography. The cerebral AV shunt created by the VGAM can increase cardiac preload and lead to congestive heart failure (CHF). Ultrasound signs of heart failure, such as cardiomegaly, triscuspid insufficiency, polyhydramnios, pericardial and pleural effusion, edema and ascites carry a poor prognosis. Antenatal fetal MR imaging will confirm the diagnosis and allow assessment of any pre-existing damage to the brain. Digital subtraction angiography (DSA, Philips) remains the gold standard technique for the evaluation of cerebral vessels, it offers precise evaluation of the VGAM angioarchitecture and provides access for endovascular treatment of the lesion. For indications of treatment, most dedicated centers use the Bicêtre Neonatal Evaluation Score (BNES) which gives information about the significant non-neurological outbreak and the neurological status. A score < 8/21 implies a decision of no treatment, a score between 8 and 12/21 requires an emergency endovascular intervention, a score > 12/21 implies the decision to manage with medical treatment until the child is at least 5-6 months old. This study has been approved by the ethical committee under protocol number P2022/256 “Prise en charge des anomalies cérébrales vasculaires à l’hopital Erasme”.

### Zebrafish husbandry

Zebrafish (*Danio rerio*) are maintained at 28°C on a 14 h light/10h dark cycle. Embryos were obtained and raised under standard conditions in accordance with European and national ethical and animal welfare guidelines (Protocol approval number: CEBEA-07 GOS IBMM). Mutant and transgenic lines used in this study are *Tg(kdrl:GFP)^s8^*^43^, *Tg(gata1a:DsRed)^sd2^, Tg(kdrl:ras-mCherry)^s896^, Tg(kdrl:NLS-mCherry)^is4^, rasa1a^ulb28^*(generated in this study) and *ephb4a^hu3445^* (obtained from ZIRC). The developmental stages were determined according to Kimmel et al^71^.

### Mutant generation

The *rasa1a* allele was generated using the CRISPR/Cas9 technology as described previously^72,73^. The targeting sgRNAs were designed using the chopchop.cbu.uib.no website and were cloned into pT7-gRNA using the following primers: rasa1a-fwd: 5’-TAGGATTGCGTCAGGCTCGGACAC-3’ and rasa1a-rv: 5’-AAACGTGTCCGAGCCTGACGCAAT-3’. The sgRNA was transcribed from the BamHI linearized pT7-gRNA vector using the MEGAshortscript T7 transcription kit (Thermo Fisher Scientific) and injected at 50 pg into one-cell stage zebrafish embryos, together with 150 pg of nls-zCas9-nls mRNA transcribed from pT3TS-zCas9-nls vector (Addgene # 46757) using the mMessage mMachine T3 Kit (Ambion). The efficiency of somatic gene disruption was assessed using high-resolution melt analysis on the Illumina Eco Real-Time PCR system and further validated by Sanger sequencing.

### Morpholinos injection and Microangiography

0.25 ng of the *tnnt2a* morpholino (CATGTTTGCTCTGATCTGACACGCA, GeneTools, Eugene, OR), 1 ng of the *rasa1a* morpholino (TAATCTCCACACATCGCAATGATCC, GeneTools, Eugene, OR), 0.5 ng of *ephb4a* (CTGGAAAACACACACGAGAGATAGA, Genetools, Eugene, OR) and 2 ng of a standard control morpholino (CCTCTTACCTCAGTTACAATTTATA Genetools, Eugene, OR) were injected into one-cell stage embryos. To assess DLV perfusion, microangiography was conducted by injecting 1 nL of 10 kDa cascade blue (10 mg/mL) into the bloodstrean through the Duct of Cuvier of anesthetized 36 hpf embryos using a micromanipulator. Embryos were mounted dorsally and imaged individually 30 min post-injection to capture tracer distribution in the vascular network.

### Phenotyping

Phenotypic analysis of *ephb4a* MO, *rasa1a* MO and control larvae was conducted at 3 dpf using a Leica M165 stereoscope to identify morphological abnormalities, including cardiac and cerebral edema. For *ephb4a* MO larvae, Z-stack imaging of the DLV was conducted using a spinning disk confocal microscope (Yokogawa CSU-X1 on a Zeiss inverted stand) with a 20X/0.5 Plan Apo objective and Metamorph software. Imaging of *rasa1a* MO was performed using a Zeiss LSM710 confocal microscope with a 20X/0.8 dry objective and driven by Zen Blue software. For mutants and WT larvae, general vasculature imaging was obtained using a Nikon AX R confocal microscope equipped with a 25X/1.05 Plan Apo Lambda S silicone objective. Large field-of-view images (3×1) were acquired using a 10X/0.45 Plan Apo Lambda D lens and analyzed using NIS software. Cardiac edema and ISV defects—characterized by vessel duplication or absence of perfusion—were quantified from large Z-stack acquisitions obtained using the 10X objective on the Nikon AX R or a Leica stereoscope. Characterization of choroidal and mural malformations was performed using Z-stack acquisitions with the same Nikon AX R confocal microscope equipped with a 25X silicone immersion objective. Choroidal malformations were described as two dissociated perfused vessels on the anterior segment with or without two or three distinctive vessels on the posterior segment. Mural malformations were characterized by vessel dilatation in the posterior segment.

### Cardiac function in zebrafish

Ventricular and atrium volumes were determined via video recordings captured using the Nikon AX confocal microscope (Nikon Instruments Inc). Imaging was performed using the 25X/1.05 Plan Apo Lambda S silicone immersion objective at an average acquisition speed of 10 frames per second. This data was calculated by extracting measurements of long-axis (lax) and short-axis (sax) ventricular and atrium diameters from static images obtained at the end of systole, exhibiting maximal dilation, and the end of diastole. NIS software was utilized to carry out these measurements, which were then used in the V = 0.5 x sax^2^ x lax formula to determine both atrium and ventricular volumes. To determine the stroke volume, we computed it by subtracting the average diastolic ventricular volume from the average systolic ventricular volume.

### Imaging and Hemodynamics parameters analysis

Embryos were imaged using a Nikon AX R confocal microscope *(Nikon Instruments Inc*) using the 10x, 20x and 25x silicon immersion objective, Nikon spinning disk microscope using the 40x immersion objective, spinning disk confocal microscope (Yokogawa CSU-X1 on Zeiss inverted stand) using a 20X/0.5 Plan Apo objective and Leica M165 stereoscope. The live imaging experiments could be done after the embryos or larvae were anesthetized with tricaine (19.2 mg/L, Sigma-Aldrich) , dechorionated and immobilized in 1% low melting point agarose in a glass-bottom Petri dish (MatTek Corporation, Ashland, MA). After imaging, the embryos were genotyped using high-resolution melt analysis (Eco Illumina real-time PCR system). Images of the DLV embryos were acquired with the Nikon AX R with a 25x objective and specifically the large image (3×1) of embryos was performed with a 10x objective. For the living zebrafish embryos immobilized in 1% low melting point agarose E3 medium and additionally supplemented with N-Phenylthiourea (30 mg/L, Sigma-Aldrich) and Tricaine (19.2 mg/L, Sigma-Aldrich). For the time-lapse analysis, we employed two approaches: the Nikon AX R confocal microscope with a stable temperature of 28.5 °C was maintained using a heating chamber, and the X-Light V3 spinning disk confocal microscope (Crest optics) with a NiR Apo 40x/0,80 DIC N2 water immersion objective, specifically for the overdose of tricaine experiment (4X). Assembly of confocal stacks and time-lapse movies was conducted using NIS software. Blood flow measurement was performed by rapidly tracking erythrocyte movement using the transgenic line *Tg(kdrl:GFP;gata1a:DsRed)*. For the frequency profile (Figure 3), we used a spinning disk confocal microscope (Yokogawa CSU-X1 on Zeiss inverted stand) equipped with a sensitive and ultra-fast EMCCD camera (Photometrics Evolve 512) and a 20X/0.5 Plan Apo Lambda D objective were used. For each sample, the displacement of erythrocytes required an acquisition speed of 200 frames per second with a number of 3000 frames. This acquisition speed allows automated analysis of the velocities by tracking the erythrocytes individually, image by image, via the scientific imaging software Fiji and the image analysis module “TrackMate”^74^. This software allows both to obtain the plot of the instantaneous velocities of erythrocytes and calculate the average velocity of these by integrating the velocity peaks. For the calculation of the fundamental frequency, the frequency profile (Figure 2) and blood parameters (Figure supp 2) , we used a spinning disk confocal microscope (X-Light V3, Crest optics) equipped with a fast camera (Prime BSI Scientific CMOS (sCMOS)) with an acquisition speed of 750 frames per second and a NiR Apo 40x/0,80 DIC N2 water immersion objective. The fundamental frequency of the red blood cells’ velocity was programmatically assessed using MATLAB version R2021b. Velocity and time data generated by “TrackMate” were imported as vectors. The sampling frequency was calculated as the inverse of the average time step between each point of the time vector. Subsequently, a frequency vector was computed using the formula f = (0:n-1)(fs/n), following the Nyquist-Shannon sampling theorem. The discrete Fourier transform of the velocity vector was then computed using the FFT function. The findpeaks function was applied to the FFT spectrum to identify the peak corresponding to the fundamental frequency. Finally, the peak position was used to determine the value of the fundamental frequency.

The average diameter measurements were performed on a maximum stack projection. We manually analyzed the vessel diameter at three different points along its length via NIS software. The data set of average globule velocities and mean diameter in each vessel allowed us to estimate the shear stress. Shear stress was estimated by measuring the average diameter of the blood vessel and the mean velocity of red blood cells as an approximation of the debit, we did not consider the non-Newtonian nature of blood at this scale, but the blood vessels in zebrafish embryos have roughly similar dimensions in the micrometer range. We used the following formula to estimate it: 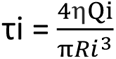 where η is the blood viscosity, Qi is the average velocity (μm.s^-1^) and Ri is the vessel diameter (µm) To demonstrate the generation of two independent streams, the tracks of red blood cells produced by “TrackMate” were sorted using a Python program (Jupyter Notebook version 6.4.5 running using conda 4.14.0 and Python 3). For each track, the initial and final positions of the red blood cells were identified with respect to the x and y axis. A central axis was then created to follow the direction of the DLV. Tracks were then sorted based on the position of their initial ad final points relative to the central axis. Tracks were then assigned different colors based on this sorting.

### iDISCO protocol for tissue transparization and volumetric imaging

The described method was adapted from the iDISCO protocol^75^. The mice tissues were collected from mice cadavers used for other purposes (protocol 652N) and fixed overnight at 4°C in 4% PFA in PBS. After fixation, the samples were washed three times for 1 hour each in PBS and subsequently dehydrated using a series of methanol/PBS solutions (50%, 80%, 100%, 100%) for 1 hour at room temperature. The samples were then bleached in 5% H2O2 in 20% DMSO/methanol overnight at 4°C. After bleaching, samples were washed in 100% methanol 1h twice, then in 20% DMSO/methanol for 2h, then in 80% methanol for 1h, 50% methanol for 1h, PBS for 1h twice, and finally in PBS/0.2% Triton X-100 for 1h twice before staining. Pretreated samples were incubated in PBS/2%TritonX-100/20%DMSO/0.3M glycine at 37°C overnight. Blocking in 2% Triton X-100/Intercept (PBS) Blocking Buffer (LI-COR) at 37°C for 2 days, the samples were washed twice for 1 hour each in PBS containing 0.1% Tween-20. Subsequently, the samples were incubated with the primary antibody PECAM-1 (at a dilution of 1:500) in 2% Triton Blocking Buffer at 37°C for 4 days. After incubation, the samples were washed with PBS containing 0.1% Tween-20 at various time intervals (10 minutes, 15minutes, 30 minutes, 1 hour, 2 hours, and overnight). Next, the samples were incubated with a secondary antibody in 2% Triton Blocking Buffer at 37°C for 4 days. Further washes were performed using PBS with 0.1% Tween-20 at various time intervals (10minutes, 15 minutes, 30 minutes, 1 hour, 2 hours, and 1 day). Immunolabeled tissues were then incubated overnight in a solution of 50% Tetrahydrofuran/H2O, followed by incubation in 80% tetrahydrofuran for 1 hour and 100% tetrahydrofuran for 2 hours. Subsequently, the samples were incubated in Dichloromethane until they sank to the bottom of the vial. Finally, the samples were cleared and stored in DiBenzyl Ether. The tissues were fixed on glass bottom dishes and imaged using a Nikon AX confocal microscope with a 25X/1.05 Plan Apo Lambda S silicone objective.

### Cell culture

HUVECs were purchased from Lonza and cultured on gelatin-coated tissue culture dishes with EGM-2 culture medium (Lonza). Cells were used between passages 3 and 5 for experiments. siRNA transfection was performed with Lipectamine RNAiMax and ONTarget siRNA (Horizon Discovery Bioscience), reference: L-005276-00-0005 for Rasa1 and D-001810-10-05 for the non-targeting scramble siRNA. VEGF 50 ng/mL (Thermo Fisher) was used as a chemical agonist. Shear stress application, western blot experiment and immunostaining of HUVECs were previously described^9^. Briefly, cells were seeded on fibronectin-coated slides (20 µg/mL) and grown to confluence. For short-term experiments (up to 30min), cells were starved in EBM-2 medium with 0.2% FBS for 4 hours. For long-term experiments (more than 4 hours), cells were straved for 1 hour in 0.2% FBS-EBM. Shear stress of 12 dynes/cm² was applied in a parallel flow chamber for the indicated times. Cells were fixed with 4% PFA, permeabilized with 1% Triton X-100, blocked, and incubated for 1 hour with Phalloidin and DAPI at room temperature. For Western blots, cells were washed with cold PBS, lysed in SL-DOC buffer, and proteins were separated via SDS-BOLT, transferred to nitrocellulose membranes, blocked with blocking buffer, and probed with primary antibodies against pERK1/2, pAKT ser473, RASA1 and actin, followed by detection with DyLight PEG-conjugated secondary antibodies.

### Construction of a millifluidic flow chamber

The chamber model was designed on Solidworks. It was then 3D printed using a Formlabs 3D Printer with Dental SG material. After printing, the chamber was filled with SYLGARD 184 Silicone and incubated at 65°C overnight to solidify. On the second day, the silicone model was carefully released from the chamber. The chamber was assembled with a glass slide coated with fibronectin and HUVECs. The flow was introduced into the model through two inputs and one output using microtubes connected to an Elveflow microfluidics pump OB1 (Elvesys), generating separate flows for each inlet with controlled pressure. Flow meters were used to control debit right before the inlet. For our experiments, we used a phase shift of 180° for the pressure wave between each inlet. The micro fluorescence beads diluted in water (FluoSpheres™ Polystyrene Microspheres, 1.0 µm, blue-green (430/465)) were perfused into the right channel, and the left channel was perfused with pure water. Images were obtained using a fast timelapse using a spinning disk microscope (X-Light V3, Crest optics) equipped with a fast camera (Prime BSI Scientific CMOS (sCMOS)) and a Plan Apo 20x/0.75 DIC N2 objective.

### Intracellular calcium imaging

HUVECs (LONZA) obtained commercially were utilized between passages 4 and 5 and cultured using EGM-2 medium. The cells were seeded onto glass coverslips coated with fibronectin at a 20 μg/mL concentration. Once the cells reached confluency, they were incubated with a calcium dye working solution in an incubator at 37°C for 40 minutes. The calcium dye working solution was prepared by mixing CalbryteTM-520 AM (AAT Bioquest) with 4% FBS-EBM to achieve a final concentration of 5 μM. Additionally, 0.04% Pluronic F-127 was included in the solution. Before imaging, the medium was changed to 2% FBS-EBM at room temperature for 10 minutes. Coverslips containing cells were embedded in the microfluidics chamber and perfused with 2% FBS-EBM. We captured images within the collecting channel at intervals of 2 seconds for 30 minutes using a spinning disk microscope (X-Light V3, Crest optics) equipped with a fast camera (Prime BSI Scientific CMOS (sCMOS)) and a Plan Apo 20x/0.75 DIC N2 objective.

### Pharmacological treatment of *rasa1a* and *ephb4a* mutants

We performed two different protocols to study the imbalance in flow activation of PI3K/ERK signaling. In the first protocol, dechorionated embryos were incubated from 36 hpf to 60 hpf (time of the beginning of the formation of the DLV) and from 60 hpf to 72 hpf in E3 medium (0.3 g/L Instant Ocean Salt, 75 mg/L CaSO_4_; 1 mg/L Methylene Blue) supplemented with PTU (30 mg/L, Sigma-Aldrich) and the different drugs DMSO (Sigma Aldrich), 1µM Selumetinib (TOCRIS), 1µM YS-49 (MCE) and a combination of this two drugs. Embryos were anesthetized with tricaine (19.2 mg/L, Sigma-Aldrich) immobilized in 1% low melting point agarose in a glass-bottom Petri dish (MatTek Corporation, Ashland, MA), mounted dorsally and individually imaged using a Nikon AX confocal microscope with a 25x silicon immersion objective. In the second protocol, dechorionated embryos are anesthetized, immobilized, and mounted dorsally are imaged with the 25x at 72 hpf. Embryos were taken away from the glass-bottom Petri dish and were incubated from 72 hpf to 96 hpf (after the complete development of the DLV) in E3 medium supplemented with PTU and the same drugs previously described. After anesthesia, embryos are remounted to image the DLV after drug treatment with the same imaging parameters. For the two protocols, after the imaging (at 72 hpf for the first one and 96 hpf for the second), the embryos were genotyped using high-resolution melt analysis (Eco Illumina real-time PCR system).

### Statistical analysis

Statistical analysis was performed using the GraphPad Prism 9 software. Data are expressed as mean ± SEM, and the distribution is displayed within the graphs. P values were calculated by various tests, depending on the normality of the data’s distribution in GraphPad Prism, as indicated in the figures’ legends.

### Antibodies table

Antibodies table:
| Antigen | Vendor | Cat. |
| --- | --- | --- |
| β-Actin | SIGMA | A5441 |
| PECAM1 | R&D | AF3628 |
| Phospho-Akt (Ser473) | Cell Signaling | 4060 |
| Phospho-ERK1/2 (Thr202/Tyr204) | Proteintech | 28733-1-AP |
| RASA1 | Invitrogen | MA4-001 |
| DAPI | Invitrogen | 62248 |
| Alexa Fluor™ Plus 647 Phalloidin | Invitrogen | A30107 |

