## Supplementary figures and images for "Reversal of vein of Galen aneurysmal malformation by stimulation of flow-mediated vessel fusion"

### Supplemental Figure 4

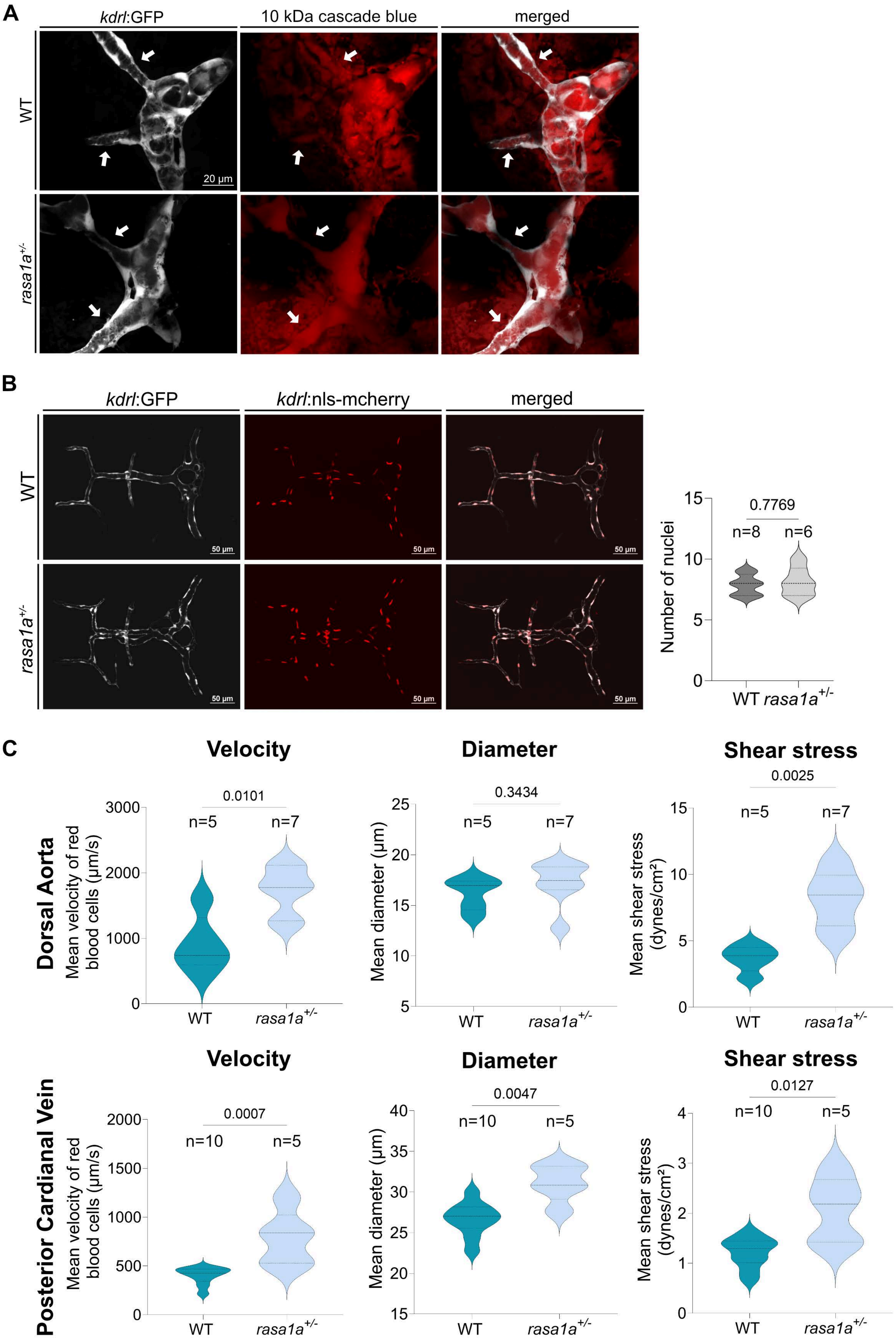
